# ObLiGaRe doxycycline Inducible (ODIn) Cas9 system driving pre-clinical drug discovery, from design to cancer treatment

**DOI:** 10.1101/2020.04.06.028803

**Authors:** Anders Lundin, Michelle J. Porritt, Himjyot Jaiswal, Frank Seeliger, Camilla Johansson, Abdel Wahad Bidar, Lukas Badertscher, Emma J. Davies, Elizabeth Hardaker, Carla P. Martins, Therese Admyre, Amir Taheri-Ghahfarokhi, Jenna Bradley, Anna Schantz, Babak Alaeimahabadi, Maryam Clausen, Xiufeng Xu, Lorenz M. Mayr, Roberto Nitsch, Mohammad Bohlooly-Y, Simon T. Barry, Marcello Maresca

## Abstract

The CRISPR-Cas9 system has increased the speed and precision of genetic editing in cells and animals. However, model generation for drug development is still expensive and time-consuming, demanding more target flexibility and faster turnaround times with high reproducibility. We have generated a tightly controlled ObLiGaRe doxycycline inducible SpCas9 (ODInCas9) transgene. Targeted ObLiGaRe resulted in functional integration into both human and mouse cells culminating in the generation of the ODInCas9 mouse. Genomic editing can be performed in cells of various tissue origins without any detectable gene editing in the absence of doxycycline. Somatic *in vivo* editing can model non-small cell lung cancer (NSCLC) adenocarcinomas, enabling treatment studies to validate the efficacy of candidate drugs. The ODInCas9 mouse can be utilized for robust and tunable genome editing allowing for flexibility, speed and uniformity at reduced cost, leading to high throughput and practical preclinical *in vivo* therapeutic testing.

## INTRODUCTION

Genetic manipulations in cells and organisms are used to understand the role of genes in a physiological or disease context. One strategy has been to exploit nucleases such as zinc finger nucleases (ZNFs), transcription activator like effector nucleases (TALENs) and clustered regularly interspaced short palindromic repeats (CRISPR) for targeted insertions and deletions (indels) or precise genome edits[1–4]. Among those nucleases, the CRISPR-Cas9 system has revolutionized precision molecular genetic approaches, accelerating the generation of genetically engineered cells and animal models (reviewed in Adli, 2018[5]).

Genetic manipulation of mice to create disease models or validate therapeutic targets are fundamental to most areas of biology. In oncology research, there is significant investment in genetically engineered mouse models (GEMMs) where gene expression is both spatially and temporally controlled to enable more faithful modelling of equivalent human tumors. While GEMMs are powerful tools, mouse strain production is both time-consuming and costly and needs to be repeated for each new engineering. Moreover, GEMMs are expensive to maintain with each specific model requiring extensive breeding to maintain a single model. A further requirement of somatic multiplex-mutagenesis is extensive mouse intercrossing for the generation of relevant experimental cohorts of multi-allelic mutant mice[6]. In addition to long breeding programs, the tumor models have a long indolence time, which often precludes the use of these models for drug testing[7]. When these models have a latency period conducive to drug testing, due to the complex nature of breeding programs required, often only small numbers of tumor bearing animals can be recruited onto study at a given time, compared to more traditional xenografted or explant cancer models[8]. To reduce cost and increase output, it is preferential to have tunable timelines to adapt tumor development to treatment time based on compound activity and mode of action[6].

In contrast, non-germline GEMMs (nGEMMS), where somatic cells carry engineered alleles but not germline cells, provide an option to shorten production time lines and increased genomic heterogeneity[9]. However, timelines to establishment of tumor models are still long, up to months before having a therapeutic window (reviewed in Heyer, 2010[10]). Moreover, traditionally used xenografted nGEMMs are also often immunodeficient, limiting its applicability[6]. Furthermore, cancer prone non-germline chimeric mice, produced by injecting genetically engineered embryonic stem cells into blastocysts of a mouse line, have an increased variability related to tumorigenesis between individual mice[11]. This negatively affects standardizing treatment studies in these models.

Instead, nGEMMs which rely on *in vivo* delivery of editing components to generate somatic genomic changes of oncogenes and/or repressor genes, provide greater flexibility in target turnover and recapitulation of tumor microenvironment, including immune effects. Gene manipulation is often achieved by viral delivery, using either an adeno-associated virus, adenovirus or lentivirus[12, 13]. Due to limitations in the viral packaging capacity, nucleases, target sequences and repair templates are often separated into different vectors, resulting in lower editing efficacy[14]. The introduction of constitutive expressing Cas9 mice, Rosa26-Cas9, reduced the number of components that needed to be delivered to the cell thereby increasing editing efficacy[13, 14].

The utility of constitutive Cas9 systems can however be limited by off-target effects and inflammatory responses if Cas9 expression is not controlled. Inducible expression systems including recombinase mediated cassette exchange (RMCE), Cre-recombinase or tetracycline (Tet) sensitive systems, have been used to generate inducible Cas9 mice[12, 13, 15–20]. Cre-systems are limited by potential genotoxicity due to cryptic recombination sites in mammalian genomes[21], which is not observed with doxycycline (dox) or Tet-inducible systems. Moreover, a Cre based system allows only for a single activation followed by constitutive expression while a Tet-inducible system can repeatedly be turned on and off, thereby avoiding downstream effects of constitutive expressions. However, tight regulation of the Tet-system is challenging so common practice is to split the two major components (Tet repressor and inducible promotor) and introduce the two different parts in different loci, thus complicating the genetic engineering process[22].

Here, we describe the generation a novel transgene, ObLiGaRe Doxycycline Inducible Cas9 (ODInCas9), an all in one, universal Tet-On system where a combination of insulators placed in a modular vector allows tight temporally regulated expression of *Streptococcus pyogenes* Cas9 (SpCas9). The ODInCas9 transgene carries human and mouse ZFN sites for targeted ObLiGaRe-mediated integration into both human and mouse genomes, enabling the generation of Cas9 inducible human iPSCs and mouse ESCs. ODInCas9 is functional and tightly controlled *in vitro* and *in vivo*. The ODInCas9 mice demonstrate inducible Cas9 expression in all tissues and can be repeatedly induced with no Cas9 detection upon dox withdrawal. Delivery of sgRNAs as a single component to Cas9 expressing cells demonstrate high editing efficacy resulting in disease phenotypic readouts with no detectable edits in the absence of dox. The ODInCas9 cassette offers oncology research the ability to introduce multiple sequential genetic alterations and better replicate the natural mutation patterns accumulated in tumors with time. Further, delivery of single stranded DNA template for increased homologous directed repair can help establish a more precise oncogene driven cancer model. The ODInCas9 mouse allows tissue-specific induction of tumors in a relevant niche that can develop and progress to late stage disease much like human tumors. The ODInCas9 mouse model provides a robust and tunable cancer induction allowing for flexibility, speed and uniformity at reduced cost, leading to high throughput and practical preclinical *in vivo* therapeutic testing models.

## RESULTS

### ODInCas9 transgene expression in multiple cellular backgrounds

To achieve inducible Cas9 expression in human and mouse cell lines of various tissue origins the novel ODInCas9 transgene cassette was developed. Inducible Cas9 expression by a tetracycline response element promoter (TRE3G) provide simple and fast genomic engineering. Inverted zinc finger nuclease (ZFN) sites targeting Rosa26 or AAVS1 of mouse or human genomes, respectively, facilitates targeted transgene integration using Obligate Ligation-Gated Recombination (ObLiGaRe)[3]. β-globulin insulators downstream of Tet-On 3G, which flanks TREG3-Cas9-T2A-GFP, allow for the generation of an all-in-one system without expression in the inactive state (Fig. 1a, b).

**Figure 1.**
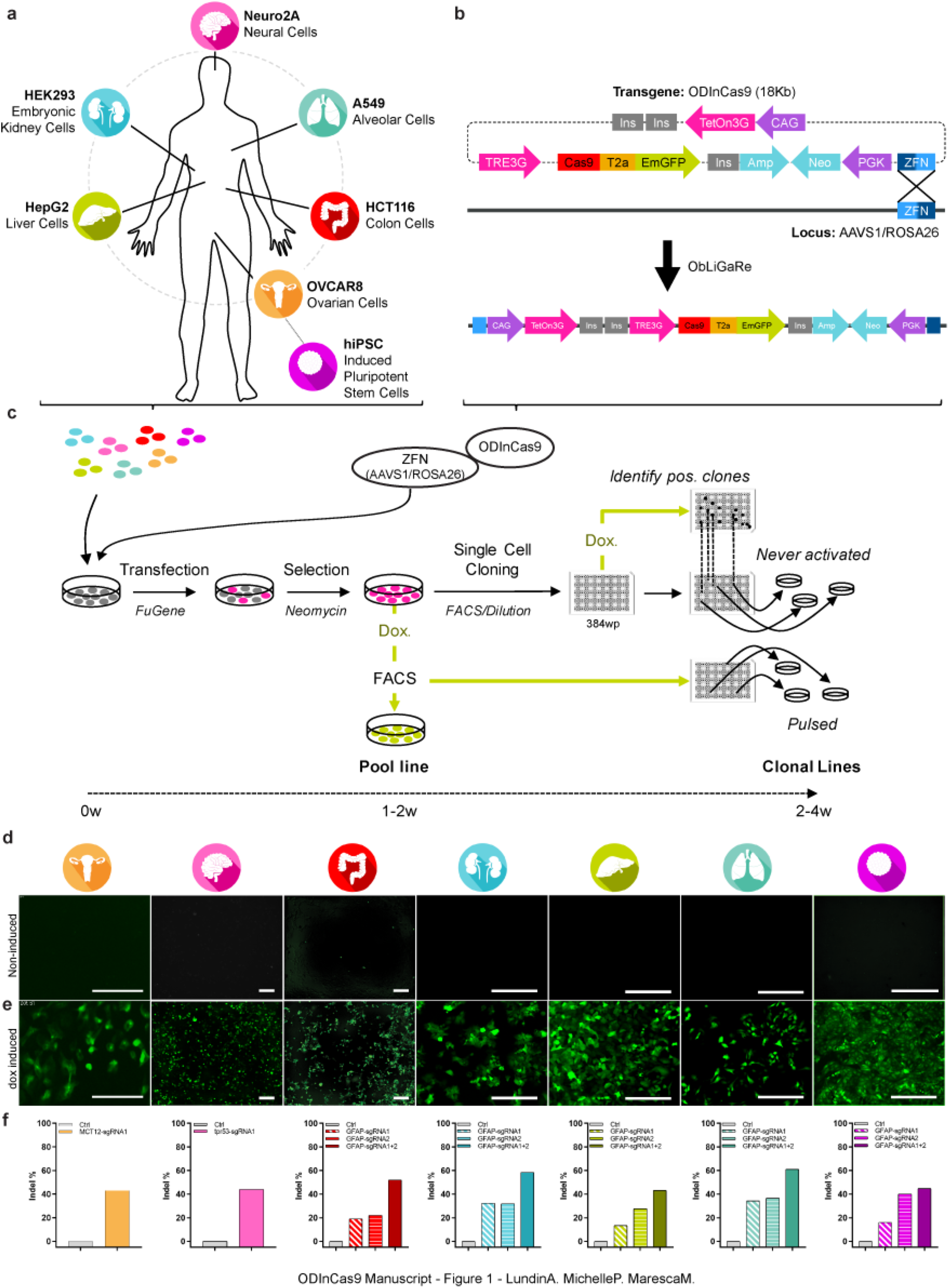
ObLiGaRe Dox Inducible (ODIn) Cas9, an all in one, universal TetOn inducible system. (**a**) Human and mouse cell lines sourced from brain (N2A), lung (A549), colon (HCT116), ovarian tissue (OVCAR8), liver (HepG2), kidney (HEK293) and human induced pluripotent stem cells (iPSC) engineered using (**b**) the ODInCas9 system targeting the AAVS1 or ROSA26 locus using ZFN directed ObLiGaRe. (**c**) Outlined methodology for rapid cell line generation of ODInCas9 pools and clonal lines encountered with or without exposure to Cas9. Evaluation of GFP expression of (**d**) non-induced and (**e**) doxycycline induced ODInCas9 cell lines. (**f**) Quantification of mismatch endonuclease assay estimating indel formation post sgRNAs transfection of induced ODInCas9 cell lines targeting; MCT1 in OVCAR8, Tpr53 in N2a and GFAP in HCT116, HEK293, HepG2, A549, and iPSC. Scale bar: 300μm.

Integration of the ODInCas9 transgene into multiple human cancer cell lines, originating from organs such as ovary (OVCAR8), colon (HCT116), kidney (HEK293), liver (HepG2), lung (A549), mouse brain cells (N2a), as well as human induced pluripotent stem cells (hiPSC) (Suppl. Fig. 1a, b), confirmed universal integration. The ODInCas9 transgene offers the ability to select clones through neomycin selection and GFP fluorescent sorting to generate cell lines within 2-6 weeks (Fig. 1c). Doxycycline (dox) treatment of ODInCas9 cell lines induced strong system activation, detectable by GFP expression. However, no activation (GFP signal) was observed in the absence of dox (Fig. 1d, e). Importantly, normal karyotype, pluripotency and differentiation capacity was maintained (Supp. Fig. 1d, f, g) in hiPSC with ODInCas9 transgene integration (Supp. Fig. 1c, f). Upon activation, ODInCas9 can generate genomic edits in hiPSC, HCT116, HEK293, HepG2, A549, OVCAR8 and N2a cells. Using either a single sgRNA or paired sgRNAs between 20-60% of cells were edited (Fig. 1f). Together, this demonstrates that ODInCas9 transgene is functional in multiple cellular backgrounds of both human and mouse origin.

### Dose and time dependent control of the ODInCas9 transgene enables high editing efficiency

ODInCas9 transgene expression is tightly controlled and both transcript and protein levels correlate to doxycycline concentration and treatment times. Dose response analysis showed detectable Cas9 and GFP expression from dox induction as low as 1-5 ng/ml in both hiPSC (Fig. 2a-c) and cancer cell lines (Supp. Fig. 2 a-d). Flow cytometry analysis showed a uniform expression of GFP in ODInCas9 hiPSC (Fig. 2d) and both human and mouse ODInCas9 cell lines (Supp. Fig. 2e). Expression levels correlated to transgene copy number integration as the hiPSC ODInCas9 homozygous clonal line displayed 2 times the level of GFP intensity and Cas9 protein expression compared to the heterozygous clone (Supp. Fig. 2f-h). Continuous stimulation with dox resulted in stable expression over time. Robust expression was detected after 6 hrs induction (Fig. 2e, f). Moreover, a 1 hr dox treatment demonstrated transient GFP and Cas9 expression going down to undetectable levels after 4-5 days (Fig. 2g-i and Supp. Fig. 2i). In summary, the ODInCas9 transgene is highly sensitive across multiple cellular backgrounds demonstrating both the flexibility of stable and transient expression.

**Figure 2.**
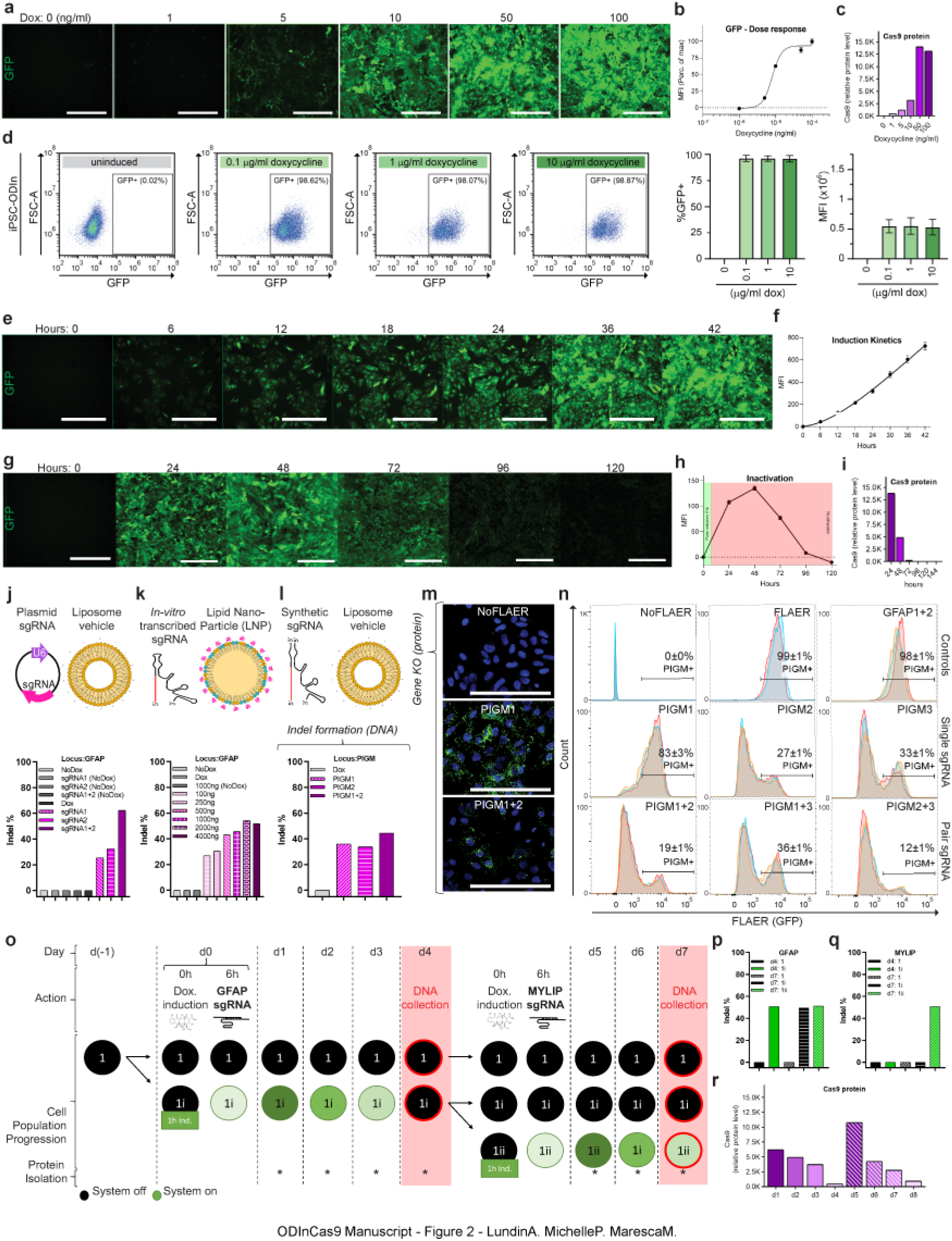
ODInCas9 system characterization and validation. (**a**) Human iPSC (hiPSC) ODInCas9 C86 GFP expression 48h post induction start displayed as (**b**) MFI evaluated by imaging (n=9 per concentration; mean ± SEM; all normalized to maximum fluorescence intensity at 100ng dox). (**c**) Cas9 protein expression in response to dox concentration from 0 to 100ng/ml (normalized to actin concentration, associated to Supplementary Fig 2c). (**d**) FACS evaluation of cell population activation at 48h post dox induction at concentrations 0.1, 1 and 10μ/ml. (**e**) GFP induction kinetics of hiPSC ODInCas9 C86 induced with 100ng/ml dox displayed as (**f**) MFI evaluated by imaging (n=9 per concentration; mean ± SEM). (**g**) Transgene expression of hiPSC ODInCas9 C86 after 1h dox treatment (10μg/ml) following washout evaluated by (**h**) GFP MFI and (**i**) Cas9 protein expression (normalized to actin concentration, associated to Supplementary Fig 2i). Endonuclease mismatch activity assay estimating Cas9 indel formation 48h post transfection of activated hiPSC ODInCas9 C86 using (**j**) sgRNA plasmid in combination with liposome vehicle delivery (associated to Supplementary Fig 2l), (**i**) in-vitro transcribed sgRNA in combination with lipid nanoparticles delivery at concentrations between 100-4000ng (associated to Supplementary Fig 2m) and (**l**) synthetic sgRNA with liposome vehicle delivery (associated to Supplementary Fig 2n). Protein knock-out evaluation of GPI anchor PIGM in hiPSC ODInCas9 C86 treated with sgRNA and paired sgRNA combinations visualized by (**m**) image representation of FLAER assay staining (**n**) and FACS analysis of PIGM+ cell population (n=2 for staining controls (NoFLAER and FLAER) and n=3 per locus targeted conditions; mean ± SEM). (**o**) Experimental design of cellular sub-culturing during repeated 1h activation of hiPSC ODInCas9 C86 at d0 and d4 with simultaneous treatment of paired sgRNAs targeting GFAP and MYLIP, respectively. Mismatch endonuclease activity assay estimating indel frequency of samples isolated at d4 and d7 at (**p**) GFAP and (associated to Supplementary Fig 2s) (**q**) MYLIP locus sites (associated to Supplementary Fig 2t). (**r**) Cas9 protein levels evaluated daily during repeated 1h activation of hiPSC ODInCas9 C86 (normalized to actin concentration, associated to Supplementary Fig 2u). *=protein isolation. MFI = mean fluorescent intensity, dox = doxycycline. Scale bar: 300μm.

The broad editing potential was demonstrated by liposomal delivery of a sgRNA targeting repetitive Alu sequences in the human genome. This revealed efficient editing causing cytotoxicity due to multitude of double stranded breaks resulting in significant reduction in both proliferation and viability (Supp. Fig. 2j-k). Liposomal and lipid nanoparticle (LNP) delivery vehicles in combination with plasmid, full length and two-component synthetic sgRNA showed targeted genomic editing (Fig. 2j-l and Supp. Fig. 2l-n). Assessing downstream effects of genomic editing guides individually targeting the surface protein GPI anchor PIGM, transmembrane protein MCT1 and essential gene CDK12 were delivered either by liposomal reagent or electroporation. Using different guides, PIGM was efficiently edited with 90% protein KO, determined by fluorescent analysis of PIGM which had been tagged using a fluorescently labeled aerolysin variant (Fig. 2m, n). Similarly, targeting CDK12, a gene involved in DNA damage response, resulted in significant reduction of cell proliferation upon protein knock-down in contrast to the non-essential transmembrane protein MCT1 (Supp. Fig. 2k, o, p). Together, ODInCas9 transgene induction in combination with standard transfection techniques of regular sgRNA components demonstrate highly efficient editing, with significant downstream effects on protein expression and cell biology.

Expression of the ODInCas9 transgene in unstimulated conditions was below the threshold of detection by different techniques. Cas9 transcript levels in inactivated hiPSC ODInCas9 were comparable to *TBX1, SOX17* and *SOX1* (Supp. Fig. 2q), biomarkers of germline differentiation, not having protein translation in pluripotent hiPSC. Doxycycline induction significantly changed *Cas9* transcript expression (Supp. Fig. 2r) to levels of *OCT4* (Supp. Fig. 2q), a highly expressed gene (Supp. Fig. 1f) driving pluripotency. Noninduced conditions have no detectable Cas9 protein or GFP intensity (Fig. 2a-i) analyzed by flow cytometry (Fig. 2d and Supp. Fig. 2i, j). Delivery of sgRNA targeting repetitive Alu elements to noninduced ODInCas9 hiPSC do not affect proliferation (Supp. Fig. 2j) and no genomic editing can be observed for delivery of sgRNA targeting specific genomic sites (Fig.2 j-l). This demonstrates that expression of the ODInCas9 transgene is prevented in the absence of doxycycline.

### Repeatable controlled induction of Cas9 expression

The ODInCas9 remains inactive after transient activation. Targeting intermediate filament, GFAP, and E3 ubiquitin-protein ligase, MYLIP, respectively, by repeating 1 hr pulse inductions at two time points, day 0 and day 4, (Fig. 2o), generated edits at each respective locus (d4:1i, d7:1ii) (Fig. 2 p, q and Supp. Fig. 2s,t) while being turned off in between editing events (Fig. 2r and Supp. Fig. 2u). No editing could be detected in the completely inactive control (d4:1, d7:1), but more importantly, the condition carrying a GFAP edit from the first induction (d4:1i) showed no edit formation at MYLIP upon repeated transfection in the inactive state (d7:1i) (Fig. 2q). All together this demonstrates that the ODInCas9 transgene is highly sensitive and versatile, additionally allowing the introduction of sequential edits.

### ODInCas9 mouse

The highly controlled features of the ODInCas9 cassette make it ideal to drive transient induction of Cas9 expression *in vivo*. As Cas9 expression is non-detectable or below the threshold of detection in the uninduced state and returns to baseline level following transient induction it provides the ideal cassette to generate a transgenic mouse without the drawbacks associated with leaky or sustained expression of Cas9. To generate a construct suitable for generation of transgenic mice the ODInCas9 transgene was inserted into the *Rosa26* locus using Rosa26 ZFN mediated ObLiGaRe[3]. Validation of transgene induction in mESC ODInCas9 cells was performed and GFP fluorescent signal and Cas9 protein expression were observed (Fig. 3a, b). The validated clone was then used to generate chimeric mice which were then expanded by breeding to a C57Bl/6NCrl background. Consistent with the tight regulation of the Cas9 expression seen *ex vivo*, heterozygote mice do not display any phenotype unless dosed with dox. Moreover, the classic mendelian inheritance observed when bred to C57BL/6NCrl mice (Fig. 3c,d) suggest no background toxicity from the construct. Homozygous mice are fertile and do not display any visible phenotype in the absence of dox.

**Figure 3.**
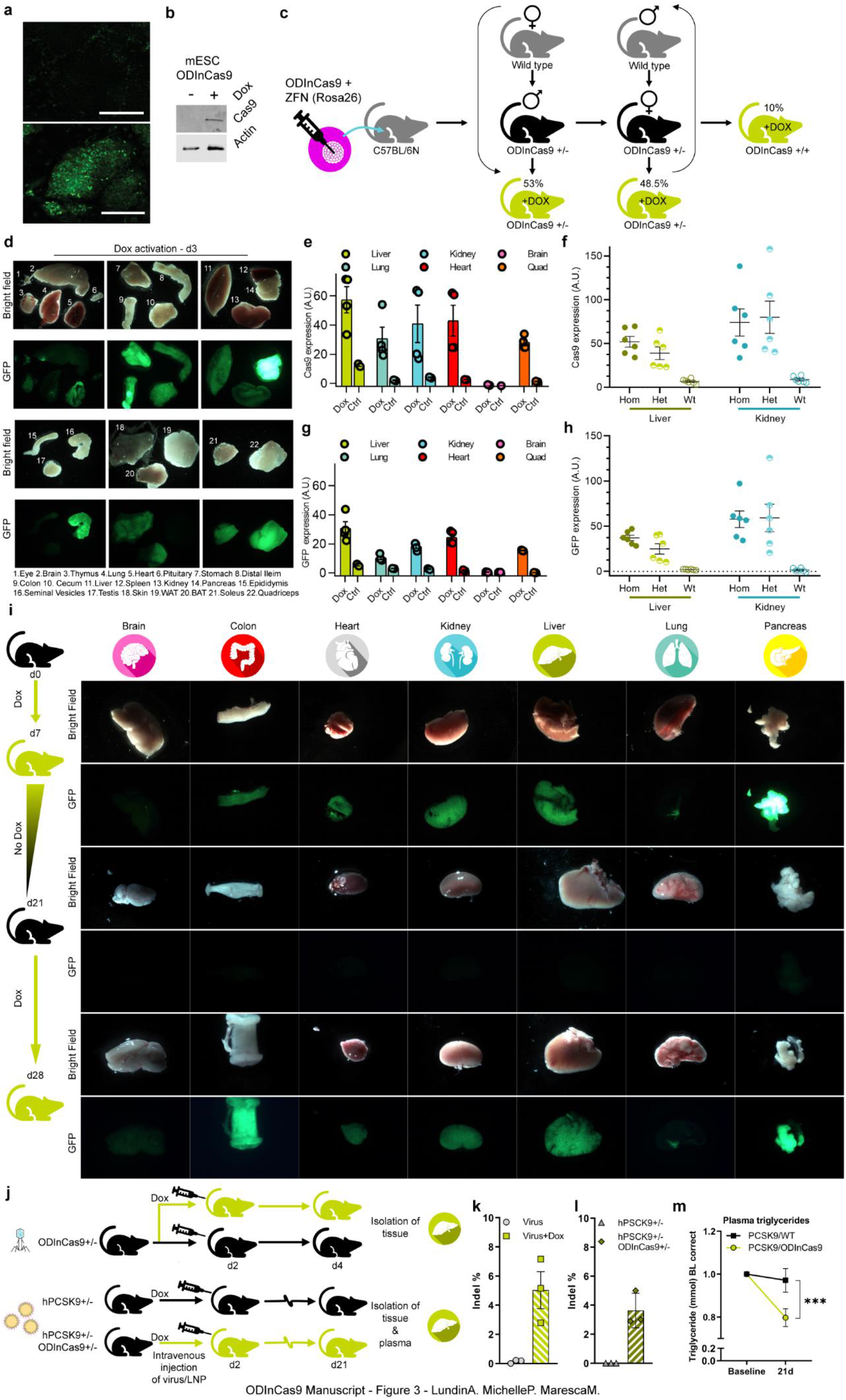
Generation and validation of the ODInCas9 mouse. (**a**) Validation of ODInCas9 mouse ESC GFP and (b) Cas9 expression with and without Dox. (**c**) Breading strategy of heterozygote and homozygote ODInCas9 mice. (**d**) Validation of GFP expression in multiple organs 72h post dox induction by fluorescent imaging. Quantification of WB of (**e**) Cas9 and (**g**) GFP protein expression in liver, lung, kidney, heart, brain and quadricep of heterozygous ODInCas9 mice (normalized to GAPDH concentration, associated to Supplementary Fig 3c). Quantification of WB of (**f**) Cas9 and (**h**) GFP expression in liver and kidney of heterozygous and homozygous mice (normalized to GAPDH concentration, associated to Supplementary Fig 3d). GFP intensity (**i**) in brain, colon, heart, kidney, liver, lung and pancreas measured at multiple time points; d7, d21 and d28 following 7d doxycycline stimulation, 14days washout without dox stimulation and 7days of dox stimulation. Schematic of (**j**) Intravenous injection of AAV targeting *Trp53* in ODInCas9 mice and LNP delivered sgRNA targeting human *PCSK9* in ODInCas9 mice crossed with the human knock in *PCSK9* mouse following dox or no dox stimulation. Indel analysis of amplicon sequencing of (**k**) *Trp53* and (**l**) *PCSK9* of isolated liver tissue. (**m**) Plasma triglycerides decreased from baseline and 21 days post LNP delivered guide in PCSK9;ODInCas9 mice (t(18)=2,45, p=0.026).

### Ubiquitous distribution of Cas9 protein in the ODInCas9 mouse

To induce Cas9 expression, animals were provided drinking water with 2 mg/ml doxycycline. Following dox induction, heterozygous ODInCas9 mice expressed Cas9 protein in all tissues, consistent with the expression pattern previously reported in the Rosa26 locus [23]. Doxycycline treatment in heterozygous mice for 24 hrs results in intense staining of the GI tract and pancreas with the lowest expression observed in skeletal muscle (Fig. 3d). 72 hrs of dox treatment increases the number of organs with high Cas9 and GFP expression; lung, liver, kidney, heart and skeletal muscle (Fig. 3e, g, Supp. Fig. 3c). Detection of Cas9 and GFP by western blotting is consistent with microscopic fluorescence (Supp. Fig. 3a, b). Homozygous ODInCas9 mice have greater Cas9 and GFP expression in the liver compared with heterozygous mice (Fig. 3f, h). No Cas9 or GFP was detected in the absence of dox stimulation (Fig. 3f, h Supp. Fig. 3a). Given that heterozygous mice give robust expression of Cas9 and to maximize animal usage from each breeding, and to minimize toxicity associated with high Cas9 induction, heterozygous mice of both sexes (aged between 10-18 weeks) were used in subsequent studies.

Broad expression of Cas9 was achieved in the heterozygous mice. Cellular distribution of Cas9 expression within organs was confirmed by immunohistochemistry (Supp. Fig. 3e-m). Cas9 is expressed in lung epithelial cells, hepatic cells in the liver, tubules around the glomerulus in the kidney and in exocrine epithelial cells of the pancreas (Supp. Fig. 3e-h). The Cas9 positive cells within the seminal vesicles were in the seminiferous epithelium in mucosal folds, epithelial cells of the intestines, skin and spleen whereas in the brain, staining was limited to ependymal cells of the lateral ventricles (Supp. Fig. 3i-m).

### Controlled repeated induction of Cas9 in the ODInCas9 mouse

Treatment of the ODInCas9 mouse with dox induces Cas9 and GFP expression and removal of dox results in loss of Cas9 expression determined via GFP fluorescence (Fig 3i). Re-introduction of dox treatment, 3 weeks after the first induction, induces Cas9 expression again (Fig 3i), demonstrating the tightness of the temporal expression of Cas9 *in vivo* in line with the observations made in cell lines.

### Targeted editing in the ODInCas9 mouse liver

To determine whether targeted gene editing in the liver could be achieved in the ODInCas9 mouse, the ODInCas9 mouse was crossed with the human *PCSK9* knock-in hypercholesterolemic mouse [24]. The liver was chosen as the target organ model as lipid nanoparticle (LNP) coated sgRNAs readily accumulate in the organ. Mice (n=18 PCSK9 Het;ODInCas9 Het and n=17 PCSK9 wildtype;ODInCas9 Het) were intravenously (i.v.) dosed with LNP containing the validated sgRNA for *PCSK9* and killed after 21 days. LNP delivery of sgRNA edited ~3.5% reads covering the *PCSK9* gene and had a physiological effect, significantly reducing plasma triglyceride levels by 20% (T-test, t(18)=2,45, p=0.026)(Fig. 3j, l, m). No editing or changes in plasma triglycerides were observed in LNP treated wild type mice.

While LNP can deliver sgRNAs to certain tissues, it is not a vehicle capable of delivering sgRNAs to tissues beyond the liver at high efficiencies. Moreover, LNP delivery is associated with inflammation and other toxicities. Virus particles are an alternative vehicle with which to deliver sgRNA with higher efficiencies and greater organ specificity. The adeno-associated virus serotype 9 (AAV9) has broad tissue tropism and is not associated with systemic inflammation. AAV9 with a sgRNA targeting *Trp53* was delivered intravenously to ODInCas9 mice. Doxycycline treated mice had significant (5%) editing in the liver after 3 days (Fig. 3j, k). In uninduced ODInCas9 mice, AAV9 delivered sgRNA resulted in no editing of the *Trp53* gene (99,9% wild type expression). This demonstrates that AAV9 delivery of sgRNA to transgenic mice bearing the ODInCas9 transgene have potential to enable targeted CRISPR modulation of multiple tissues.

### Generation of genetically complex tumor models in the ODInCas9 mouse

The ODInCas9 mouse has significant potential as a universal background in which to rapidly generate stable gene modification of tissues. Modelling tumor biology in mouse models requires tissue-specific mutation, deletion or truncation of multiple genes. Moreover, to understand the impact of a novel gene on tumor development requires time for generation of a new mouse or establishing complex breeding programs. Through conventional routes this requires long breeding programs and can be hampered by long indolence time for tumors to appear. To establish whether the ODInCas9 mouse can be used to induce key cancer mutations in a specific organ that leads to tumor formation, we delivered AAV to the lung.

As AAV9 has high trophism in lung tissue and with precedence for transforming lung tissue with inhaled viruses [12], the ability to generate lung tumors was assessed. To induce lung tumors AAV9 virus bearing sgRNA to modify the *Kras* oncogene and *Trp53* tumor suppressor were generated. A common strategy was to generate a gain of function point mutation in *Kras* (G12D or G12C) by homology directed repair and targeted editing in either the tumor suppressor gene *Trp53* (*KrasG12D;Trp53^-/-^, KrasG12C;Trp53^-/-^*) or protein kinase *Stk11* (*KrasG12D;Stk11^-/-^*) (Fig. 4a).

**Figure 4.**
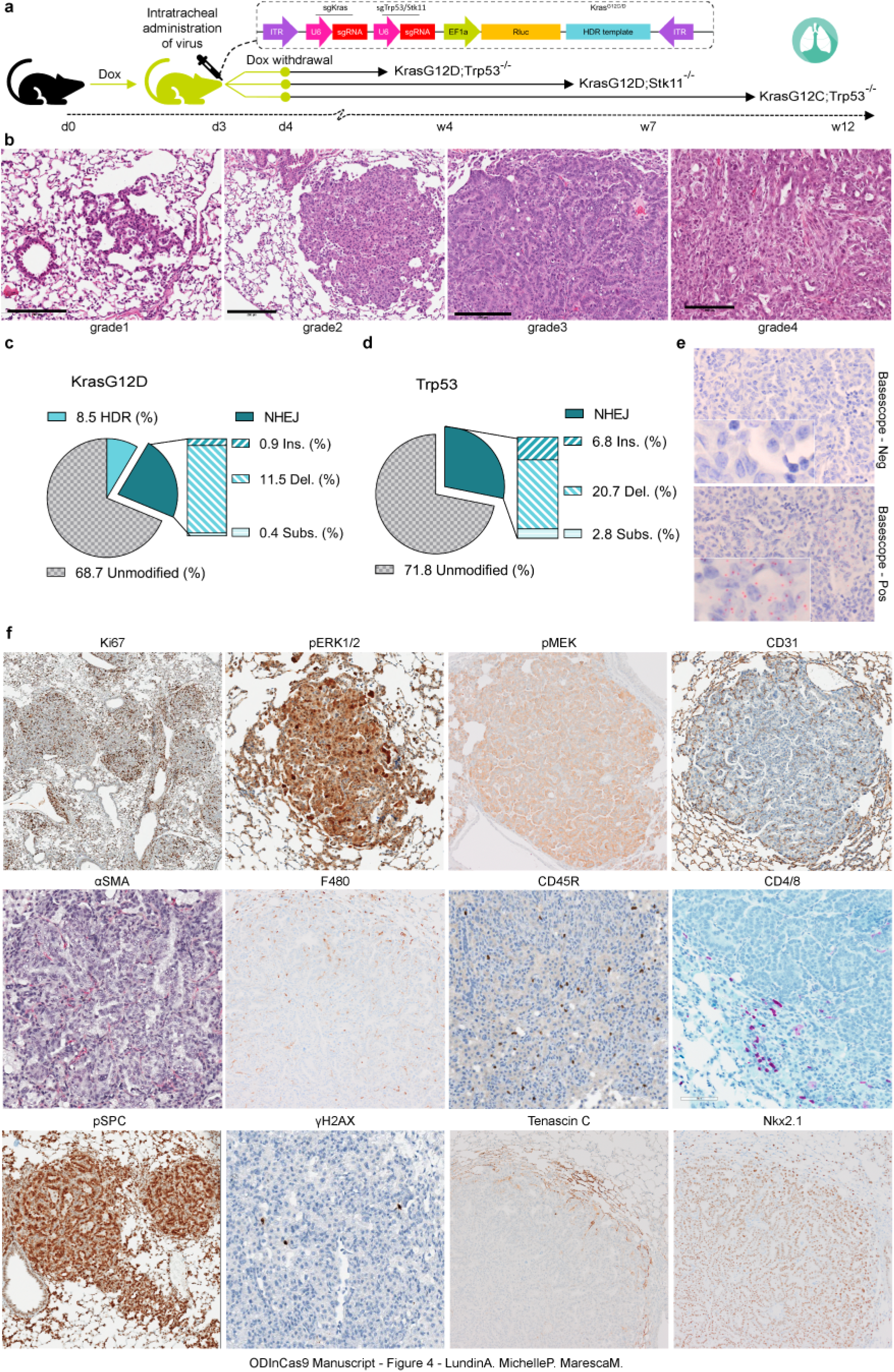
ODInCas9 mouse non small cell lung cancer models. (**a**) Schematic representation of generation of ODInCas9 *KrasG12D;Trp53^-/-^, KrasG12D;Stk11^-/-^* and *KrasG12C;Trp53^-/-^* non-small cell lung cancer models. ODInCas9 mice are treated with doxycycline 3 days before oral aspiration administration of model specific AAV targeting either *Kras* by sgRNA and HDR and *Trp53* or *Stk11* by sgRNA. Doxycycline treatment is kept 1day post viral administration. Lung adenocarcinoma kinetics is specific for each lung cancer model. (**b**) Representative HE staining of lung histology slides capturing various stages of lung adenocarcinoma (**c,d**) Classification of amplicon sequencing of lung cancer tissue at *Kras* and *Trp53* locus (**e**) DNA *in situ* hybridization targeting KrasG12D (**f**) Immunohistochemistry section of lung tumors staining for Ki67, pERK, pMEK, CD31, αSMA, F480, CD45R, CD4/8, pSPC, ɣH2AX, Tenascin C and Nkx2.1.

As described above mice were induced with 2 mg/kg dox in the drinking water, and AAV9 carrying sgRNAs were delivered i.v. or by oropharyngeal aspiration to the lung. All animals that received AAV9 developed adenomas and adenocarcinomas in the lung. Oropharyngeal aspiration restricted tumor development to the lungs. Intravenous dosing is a potential method of induction however mice dosed i.v. have fewer lung tumors compared to aspiration dosing. Further, i.v. viral dosing also results in unintended tumors in non-target tissues. We have observed a soft tissue sarcoma present in the nose skin and a cholangiocarcinoma in the liver after i.v. dosing.

A robust tumor development was observed in all models; *KrasG12D;Trp53^-/-^* (Fig. 4b), *KrasG12C;Trp53^-/-^* (Supp. Fig 4A b) *and KrasG12D;Stk11^-/-^* (Supp. Fig 4B b) each with slightly different attributes (Table 1). A key feature of all models is that they are adjustable, the tumor burden can be altered by the viral titer dosed (Supp. Fig. 4B e (n= 30)).

**Table 1:** Non-small cell lung cancer (NSCLC) models developed by AAV9 transduction in the ODInCas9 mouse

| Genetic Alterations | AAV titer (GC) | Latency to treatment | No. & Size of tumor nodes | Features relevant to model |
| --- | --- | --- | --- | --- |
| *KrasG12D;Trp53<sup>-/-</sup>* | 1 X 10<sup>11</sup> | 4 weeks | Numerous, small | Multiple tumor nodules from 3 weeks post induction, random distribution of nodules of all grades throughout the lobes (Fig. 4b). |
| *KrasG12D;Stk11<sup>-/-</sup>* | 1 X 10<sup>10</sup> | 7 weeks | Few, large | Atypical hyperplasia's, adenomas and adenocarcinomas at 4 weeks. Fewer tumors per lung lobe with rapid growth of tumors between 12- and 16-weeks post induction (Supp. Fig. 4A). |
| *KrasG12C;Trp53<sup>-/-</sup>* | 1 X 10<sup>11</sup> | 12 weeks | Numerous, small | Similar to the *KrasG12D;Trp53<sup>-/-</sup>* model however these numerous tumor foci progress at a slower rate of growth (Supp. Fig. 4B). |

### ODInCas9 NSCLC models recapitulate clinical tumor pathology

The tumors generated in the ODInCas9 NSCLC models reflect pathological features of human NSCLC. Multiple tumors were detected in each animal (Fig. 4b, Supp. Fig. 4a, Supp. Fig. 4b). Tumors increase in size over time and as early as 3 weeks post viral infection multiple grade I and II bronchial alveolar adenomas were present, progressing to grade III lung adenocarcinomas and grade IV adenocarcinomas (Fig. 4b, Supp. Fig. 4a, Supp. Fig. 4b). Commonly, adenocarcinomas are observed infiltrating through the pleura and into the mediastinum. Blood and lymphatic vessel invasion were correlated to distant metastases, primarily to gatekeeper lymph nodes. Tumor and non-neoplastic cells were highly proliferative, by means of Ki67 immunohistochemistry (Fig 4f, Supp. Fig.4A f, Supp. Fig.4B e).

Tumor cells expressed pro-surfactant protein C (proSpC) indicative of alveolar type 2 cells as the primary edited cell type (Fig. 4f). All tumors are highly vascularized (Fig. 4, Suppl. Fig.4A f, Supp. Fig. 4B e) and showed a thin aSMA positive stroma. Occasionally, low grade adenocarcinomas (grade IV) exhibited focal random expression of Tenascin C in tumor cells and extracellular matrix which was reversely correlated to the loss of Nkx2.1 in these tumor cells indicative of tumor progression (Fig. 4f).

The immunophenotype of the tumor models was characterized by the presence of all relevant major immune cell lines in the tumor or on the surface area in the adjacent lung tissue. Most malignant tumors were macrophage rich with a robust accumulation of F4/80 positive macrophages in the extracellular matrix, while the benign phenotypes show a varying number of B cells (CD45R) (Fig. 4f, Supp. Fig. 4A. Supp Fig. 4B e). CD4 T cells and CD8 cytotoxic T cells were observed in low numbers in the tumor stroma (Fig. 4f, Supp. Fig. 4A. Supp Fig. 4B e).

Eight weeks after viral administration, genomic DNA was extracted from tumor bearing lungs and targeted sequencing performed using Illumina MiSeq. In the *Kras G12D;Trp53^-/-^* model, editing of the *Kras* gene resulting in the oncogenic *Kras* form G12D, present in 8% of the sequencing reads 28% of the reads had a mutation in the *Trp53* gene (Fig. 4d). In the *Kras G12D;Stk11^-/-^* model, 7 weeks post induction, editing was 7% for *Kras* and 28% for *Stk11* (Supp. Fig. 4A c, d). In the *Kras G12C;Trp53^-/-^* model, editing was 13% for *Kras* and 28% for *Trp53* (Supp. Fig. 4B b, c). The correct insertion of each specific *Kras* point mutation G12D or G12C; was further confirmed by single point mutation sensitive *in situ* hybridization (BaseScope^®^, ACD) [25] (Fig. 4e, Supp Fig. 4B d). Further, the tumors, including atypical hyperplasia’s express pERK1/2 and pMEK confirming activation of the RAS/MAPK pathway (Fig. 4e, Supp. Fig. 4A. Supp Fig. 4B e) and correct genome editing.

### Pharmacological interrogation of signaling pathways in ODInCas9 mouse derived tumor models

To demonstrate the utility of the ODInCas9 mouse NSCLC models for preclinical testing, efficacy studies based on previous clinical [26] and preclinical studies [27] were performed (Fig. 5, Supp. Fig. 5). ODInCas9 mice (n=17) were induced and transduced by AAV9 carrying the sgRNA and homology repair template. *KrasG12D;Trp53^-/-^* tumors were allowed to progress for 4 weeks prior to treatment with vehicle (n=6) or combination chemotherapy Docetaxel and the MEK inhibitor AZD6244 (Selumetinib) (n=6) for 2 weeks (Fig. 5a). At 4 weeks there was significant tumor burden as determined by histological assessment of satellite mice sacrificed prior to treatment (Fig. 5a, e). Combination therapy resulted in the stasis of tumor growth, measured by histology at termination, preventing the 3-fold increase observed in vehicle treated animals (one-way ANOVA, F (3, 17)= 5.449, p=0.0279) (Fig. 5b, d, e). The majority of tumors responded to combination therapy, particularly the inhibition of the RAS/MAPK pathway via inhibition of MEK1/2 as demonstrated by decreased pERK staining. Further, measurement of tumor volume by MRI confirmed stasis of tumor progression with a significant treatment effect (oneway ANOVA, F (1, 19)= 5.475, p=0.0304) (Fig. 5c).

**Figure 5.**
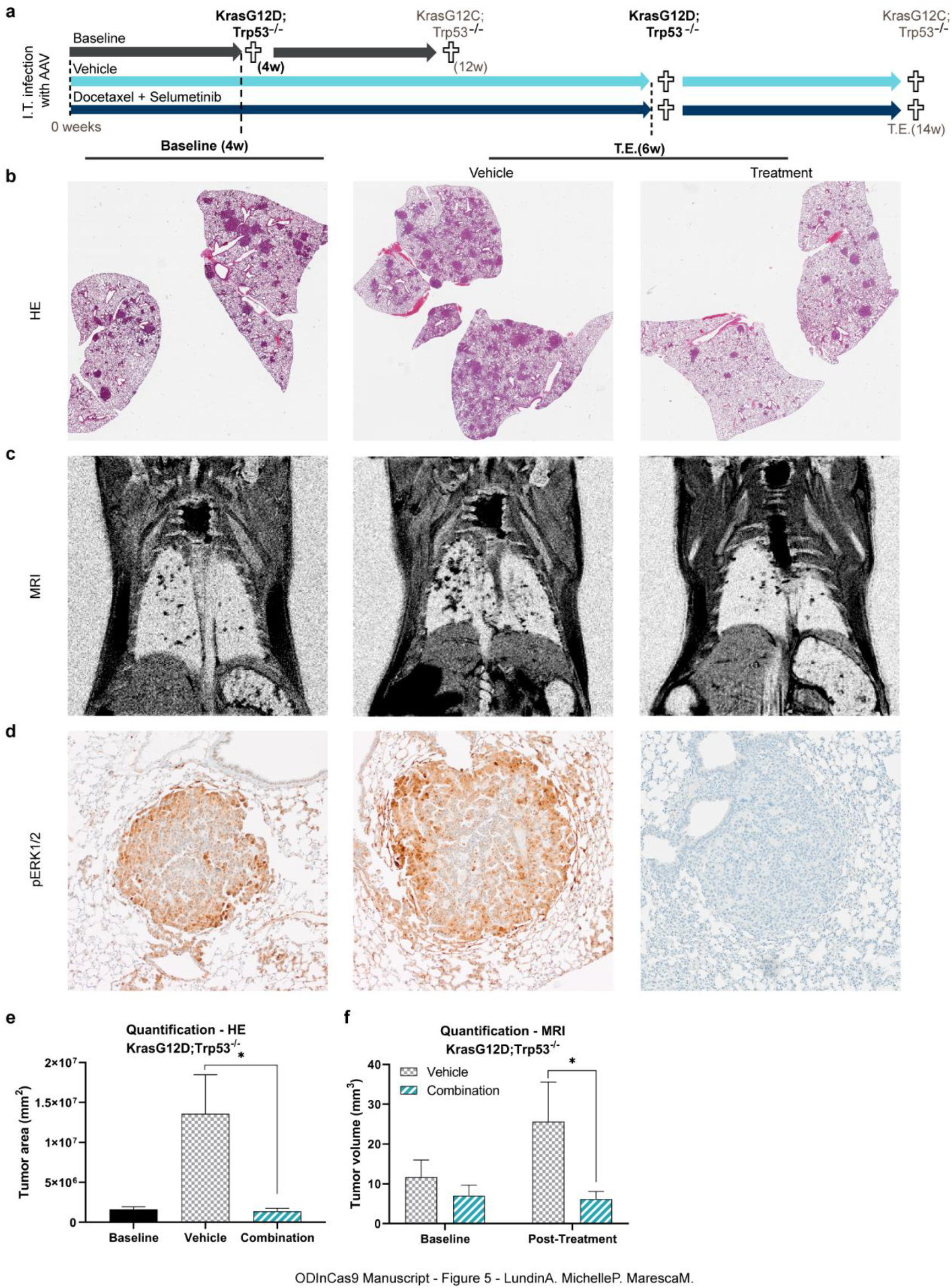
Pre-clinical efficacy study in ODInCas9 mouse non-small cell lung cancer model. (**a**) Timelines for treatment studies. ODInCas9 mice were induced with dox, dosed with AAV and allocated to treatment arm at 4w *KrasG12D;Trp53^-/-^* and 12w *KrasG12C;Trp53^-/-^*, when significant tumor burden is observed. *KrasG12D;Trp53^-/-^* mice were either killed at 4w for baseline tumor burden or treated for 4w with either vehicle or Docetaxel + Selumetinib combination treatment. Evaluation at baseline and treatment end (T.E) by (**b**) lung tissue histology (HE) (**c**) MRI (**d**) pERK1/2 immunohistochemical staining. Quantification of tumor burden at baseline, vehicle and combinatory treatment using (**e**) HE histology (ANOVA, F (3, 17)= 5.449, p=0.0279) and (**f**) MRI (ANOVA, F (1, 19)= 5.475, p=0.0304)

ODInCas9 mice (n=22) were dox induced and transduced by AAV9. Twelve weeks after *KrasG12C;Trp53^-/-^* induction mice were randomized to treatment arm and treated with either vehicle (n=8) or combination chemotherapy Docetaxel and MEK inhibitor AZD6244 (Selumetinib) (n=8) for 4 weeks. Combination treatment resulted in stasis of tumor progression and significantly decreased the percentage of lung occupied by tumor (one-way ANOVA, F (2,9)= 6.490, p=0.0180) (Suppl Fig. 5).

These results demonstrate that the ODInCas9 mouse can be used to generate genetically complex models. In this instance we have with CRISPR-Cas9 editing created both an activating mutation in *Kras* and loss of function in Trp53. Further, these models have tumors that are clinically relevant, developed in a timescale conducive to therapeutic intervention, and they are susceptible to pathway blockade and chemotherapy.

## DISCUSSION

When using Cas9 mediated gene editing to generate genetically engineered models, constitutive or leaky expression of Cas9 can have a negative impact on the utility of the model. Since short induced expression of Cas9 is important to limit off target gene editing [28, 29], our aim was to create a sensitive but tightly regulated all-in-one inducible Cas9 system that could be used in a variety of genomic backgrounds of human and mouse origins as well as generating an inducible mouse.

As the ObLiGaRe dox Inducible (ODIn) Cas9 (ODInCas9) system is an all-in-one system it allows for rapid cell line generation by a single transgene integration event compared to split Tet-On element systems[30]. In contrast to some previous all-in-one system designs [30] the ODInCas9 system shows tight and strong activation due to the insulator design allowing for the use of a strong CAG promoter without leakage. Moreover, in comparison to other inducible systems [31], the use of ObLiGaRe allowed for targeted integration of the large transgene (ODInCas9 cassette) by ZFN homologies in both human and mouse genomic backgrounds, in parallel.

Our results indicate that the ODInCas9 system is tightly regulated with no leakage detected in the cell lines or, in the mouse. In all methodologies used to investigate Cas9 expression without dox induction, non-induced cells and mouse tissue (not exposed to dox) was always at the lowest detectable limits of the experimental system used. The ODInCas9 system is very sensitive (1-5 ng/μl dox activation) and tunable. Moreover, as the system turns off upon withdrawal of dox it can be repeatedly activated for sequential editing of multiple genes, but more importantly inactivation minimizes unwanted downstream effects of the system and Cas9 exposure enabling purer efficacy studies. This emphasizes the advantage of the ODInCas9 inducible model over Cre activated Cas9 inducible systems that are constitutively active upon recombination [17, 18, 32, 33]. Tight regulation of Cas9 expression in the ODInCas9 system overcomes the leaky nature of Cre or Cas9 reported in other models [17]. Further, the Cre-loxP system is limited by the presence of cryptic loxP sites in the mammalian genome, thus genotoxicity is induced by Cre recombinase [21, 34] which is not observed with the doxycycline Tet-inducible systems

As the ODInCas9 transgenic mouse shows tight control of Cas9 expression, it offers a new approach to generate novel tumor models through direct modification of multiple oncogenes. Focusing on NSCLC, it has been possible to rapidly generate three genetically distinct tumor models, by specific modification of *Kras* G12D and *Kras* G12C, in conjunction with either *Trp53* or *Stk11*. Each of the models generated tumors with a latency of 4-12 weeks and within the course of tumor experiments, health status of the animal was only impacted by disease burden and not by non-specific side effects of the Cas9 system.

We demonstrate the successful generation of 3 models of non-small cell lung cancer. Histological and immunohistological assessment demonstrated development of multiple tumor nodes in the lungs. Tumor formation in combination with efficacy studies, including treatment with pathway specific inhibitors, significantly decreasing tumor burden suggests that the ODInCas9 mouse is a suitable replacement for traditional GEMMs. Further, transgenic mice bearing the ODInCas9 transgene can be used to generate genotype specific models at scale and in a coordinated time frame, enabling time-controlled experiments to be performed with many animals in an individual induction cohort. Ideally these complex and long-term *in vivo* experiments should edit the genome within a specific window to govern the time frame of induction of the genomic manipulations.

Unlike other tumor GEMMs, the ODInCas9 model enables development of multiple tumor models, with genetically modified alleles, from a single mouse strain. As such, this allows for a significant reduction in the time and number of animals required for the generation of each tumor model. Further, NSCLC tumor development in the ODInCas9 mouse is faster than traditional GEMMs. In some ODInCas9 models, tumors, including adenocarcinomas, can be found ~4 weeks after induction. This allows for treatment studies to be initiated at that time, rather than months after induction, as with previous lung GEMMs that targeted the same loci [7, 35–37]. Importantly, tumor burden can be easily controlled to match ideal treatment schedules through regulation of the number of infective viral titer administered.

Analysis of individual lesions revealed heterogeneous mutation patterns within the *Kras* allele. There was between 8-14% correct incorporation of the homology arm into the cut site as measured by Next-Generation Sequencing. The mRNA for each of the point mutations (*Kras* G12D and *Kras* G12C) was expressed in approximately 50% of cells within an adenocarcinoma carrying at least 1 copy of the gene. The difference in the indolence of each of the models, *Kras* G12D vs G12C and the *Trp53* with *Stk11* reveals a heterogeneity that has been reported by others [13, 17, 20] and likely results from differential usage of CRISPR-Cas9 double strand break repair mechanisms, homology directed repair, or non-homologous end joining, followed by different selective pressures over the heterogenous population of genetic mutations that arise in each model. This heterogeneity is unlike the results reported for other GEMMs and is an advantage of the inducible Cas9 systems [38]. In contrast to previous reports [39] where disruption of *Trp53* prevents the DNA damage response and increases the rate of homologous recombination, the ODInCas9 system had a similar degree of genome modification by homology directed repair to *Kras* with either *Stk11* or *Trp53* editing.

In summary, the ODInCas9 system offers a fast, reproducible and tunable genetic engineering development platform, containing a tightly controlled inducible Cas9. This model can support the preclinical pipeline with fast cell line generation (2 weeks) and *in vivo* efficacy end points as one mouse enables the development of multiple models. Further, the ability to perform multiple rounds of modification and the potential for *in vitro, ex vivo* and *in vivo* modification add to the flexibility of the ODInCas9 system enabling quick KO generation for target validation and CRISPR screening. The ability to precisely edit at different time intervals allows for preclinical replication of complex tumor mutational burden observed in oncology. Moreover, this system will decrease cost, time and reduce animal numbers compared to more traditional GEMMs as genetic manipulation of multiple loci can be achieved from a single delivery of sgRNA and homology template.

## METHODS

### Plasmids

Sequences of both the Dual AAVS1 ZFN and the ODInCas9-construct (containing *Streptococcus pyogenes SpCas9*) are provided in the supplementary information.

### Cell Culture

HEK293, HCT116, HepG2, OVACAR8, A549 and Neuro2A were all purchased from ATCC (Manassas, VA, USA) and maintained in DMEM High Glucose supplemented with 10% FBS, NEAA (1:100), and 1% penicillin–streptomycin (all from Invitrogen, Carlsbad, CA). HCT116, OVACAR8, A549 and Neuro2A were grown on tissue culture treated plates and HEK293 and HepG2, on poly-L-ornithine (20μg/cm^2^, Sigma, St. Louise, MO) coated plates. Human induced pluripotent stem cells (hiPSC) were generated as previously described [40] and maintained in a feeder free human pluripotency culturing system, Cellartis DEF-CS 500 (Takara, Japan), according to manufacturer’s instructions. All cell lines were cultured at 37 °C and 5% CO_2_ and routinely passage when reaching 80% confluency.

### ODInCas9 Cell Line Generation

#### Transfection

For ODInCas9 cell line generation of HEK293, HCT116, HepG2, Neuro2A, A549, OVCAR8 and hiPSC transfection was performed at 80% confluency. Total DNA of 500ng of ODInCas9 plasmid together with ZFN-AAVS1 plasmid at a ratio of 1:10 was mixed and transfected by lipid-based transfection using FuGene HD (Promega, Madison, WI) according to manufacturer’s instructions.

#### Transgene integration selection

For transgene integration selection HEK293, HCT116, HepG2, Neuro2A transfected cells were selected with G418 (Sigma) at a concentration of 100μg/ml starting 5 days after transfection for the following 7-9 days before single cell sorted using BD FACSAria II (BD Bioscience, San Jose, CA) for clonal cell line generation. For transgene integration selection of A549 and OVCAR8 cells were Doxycycline (Dox) (100ng/ml) treated 24h before FACS purification of GFP expressing cells for the generation of ODInCas9 pools. A549 clonal line was generated by subsequent single cell dilution (0.75cells/w) into 384wp after FACS purification. For transgene integration selection of hiPSC the cells were selected with G418 (Sigma) at a concentration of 50μg/ml starting 5 days after transfection for the following 9 days. Human iPSC clonal generation was performed by single cell dilution (0.75cells/w) into 384wp.

#### Cell line validation

ODInCas9 HEK293, HCT116, HepG2, Neuro2A, A549, and OVCAR8 cell line validation was performed by 24-48h Dox induction at 10μg/ml following GFP detection by microscopic measurement using ImageXpress Micro XLS Widefield Microscope (Molecular Devices, Sunnyvale, CA) or Incucyte Zoom (Essenbioscience, Michigan, US). Validation of transgene editing function across the ODInCas9 cells lines HEK293-C62, HCT116-12.1, HepG2-Pool, Neuro2A-C16, A549-C10, OVCAR8-Pool and hiPSC-C86 was performed by lipid-based transfection (FuGene HD) of sgRNAs expressing plasmids targeting MCT1 for OVCAR8, p53 for N2A and GFAP for HEK293, HCT116, HepG2, A549. Cells were Dox (10μg/ml) treated 12-24h before transfection followed by continuous Dox treatment until end point 48h post transfection.

For generation of non-activated ODInCas9 hiPSC cell lines confluent clonal 384wells were split into three subsequent plates for validation of transgene activation, transgene AAVS1 locus integration, transgene copy number integration and continues sub-culturing. Validation of transgene activation cells were treated with Dox (10μg/ml) for 24-48h following GFP detection using ImageXpress Micro XLS Widefield Microscope (Molecular Devices, Sunnyvale, CA). Validation of transgene locus insertion was validated by junction and AAVS1 locus PCR.

Transgene copy number was evaluated by a digital droplet PCR assay (ddPCR) purchased from BioRAD targeting the SpCas9 sequence of the ODInCas9 plasmid and normalized to a reference probe AP3B1. Digital-droplet PCR was performed according to manufacturer’s instructions (Bio-Rad, Hercules, CA). Lipid droplets are generated in part from a 20μl PCR reaction containing 100ng genomic DNA including Cas9 primers and probes. The lipid droplets undergo a cycling PCR program before analyzed using a droplet reader (QX200, BioRAD). The number of positive droplets generated by the Cas9 ddPCR assay is normalized to a reference probe AP3B1 to assess transgene copy number.

Quality of the ODInCas9 hiPSC clones was assessed by karyotyping, pluripotency marker expression and differentiation capacity. Karyotyping was evaluated by g-banding using Cell Guidance Systems karyotyping analysis service (Cell Guidance Systems, Cambridge, UK). Cell fixation was done according to instruction by Cell Guidance Systems and then shipped for analysis.

Analysis of pluripotency marker expression of hiPSC ODInCas9 clones was performed using BD Stemflow kit (BD Bioscience) according to manufacturer’s instruction. In short, cells were detached using TryPLE and resuspended in before PBS before cell fixation using Cytofix (BD). Cells were permeabilized and stained with Alexa Fluor^®^ 647 Mouse anti-SSEA-4 (clone: MC813) and PerCP-Cy™5.5 Mouse anti-Oct3/4 (clone: Clone: 40/Oct-3) including isotype controls Alexa Fluor^®^ 647 Mouse IgG3, κ Isotype Control (Clone: J606) and PerCP-Cy5.5 Mouse IgG1, κ Isotype Control (Clone: X40). Stained cells were resuspended in FACS buffer (PBSS + 2% FBS) and analyzed by FACS using BD LSRFortessa (BD Bioscience).

Differentiation capacity was assessed by applying STEMdiff™ Trilineage Differentiation Kit according to manufacturer’s instructions. In short, iPSCs were adapted from DEF-CS culturing system to mTeSR™1 (STEMcell Technologies) culturing system by seeding 40K/cm^2^ on Geltrex (ThermoFisher) for 4 days before trilineage differentiation start. Cells were seeded into three different conditions: ectoderm, mesoderm and endoderm at 200 000/cm^2^, 50 000/cm^2^ and 200 000cells/cm^2^, respectively. One day post passage cultures where switch to respective differentiation media and cultured 5days for mesoderm and endoderm lineage differentiation or 7 days for ectoderm lineage differentiation. Medium was changed daily. Cells were fixed and stained for SOX1 (AF3369, R&D Systems), SOX17 (562205, BD Bioscience) and Brachyury (X1AO2, eBioscience) assessing ectoderm, mesoderm and endoderm differentiation capacity, respectively.

#### ODInCas9 Transgene Induction Evaluation

Induction titration studies of the ODInCas9 cell lines; HEK293-C12.1, A549-C10, HepG2-Pool, hiPSC C25/C86 ranged from 1pg/ml to 100ng/ml and were applied for 24h before endpoint assessment at 48h looking at either GFP expression and/or Cas9 protein concentrations. Activation time course study of the hiPSC ODInCas9 C86 clonal line applied 50ng/ml continuously for 48h during live cell imaging. Inactivation time course study of the ODInCas9 construct in hiPSC ODInCas9 C86 clonal line was performed by a pulse induction of 10μg/ml for 1h followed by 3x PBS wash where cells were imaged and collected every day for GFP detection and Cas9 protein for 7 days.

Evaluation of cell population activation of ODInCas9 lines HEK293-C12.1 and N2A-C16 was performed 48h post Dox induction (0, 0.1μg/ml, 1μg/ml and 10μg/ml). Cells were washed with PBS, detached in 2 mM EDTA (in 1x PBS) and resuspended in FACS buffer (2 mM EDTA, 2% FBS, 1x PBS). Cells were transferred in 96-well plates (Greiner #651201) and subjected to flow cytometry on an iQue Screener PLUS (Intellicyt). GFP positive cells were identified using a BL1 detector (530 nm) and data was analyzed with the ForeCyt software.

Evaluation of transgene transcriptional activation in ODInCas9 C25/C86 lines was performed by collecting RNA 48h post experimental start of doxycycline treated (10μg/ml) and non-treated cells. Total RNA was isolated using miRNeasy Kit according to the manufacturer’s instructions (Qiagen). The quality of the RNA was assessed by a Fragment Analyzer (Advanced Analytical Technologies, Ankeny, IA). Samples with RNA integrity number >9 were used for library preparation. One microgram of total RNA was used for long RNA library construction. Illumina TrueSeq Stranded mRNA LT Sample Prep Kit (Illumina, San Diego, CA) was used to construct poly(A) selected paired-end sequencing libraries according to TrueSeq Stranded mRNA Sample Preparation Guide (Illumina). All libraries were quantified with the Fragment Analyzer (Advanced Analytical Technologies), pooled and quantified with Qubit Fluorometer (Invitrogen) and sequenced using Illumina NextSeq 500 sequencer (Illumina). Three biological replicates were sequenced per sample. For RNAseq data analysis Salmon [41] were used to quantify transcript read counts. A hybrid reference transcript was generated using Ensemble v94 and Tet-On3G and Cas9 sequences used in the plasmid. Transcript level data were collapsed to the gene level and then normalized using tximport[42].

#### Evaluation of sgRNA delivery

Single gRNAs were transfected either as plasmid, *in vitro* transcribed or synthetic. Induction with Dox (100ng/ml) was perform 12-24h before transfection and maintained to end of experiment, 48h post transfection. Cells were grown to 80% confluency and while passaged mixed with transfection reagents before seeding. End point experimental readout was to evaluate indel (see indel assessment method section) and/or protein expression (see western blot method section). Plasmid transfection was performed using FuGene according to manufacturer’s instructions transfecting 500ng plasmid per 24w having a reagent:DNA ration of 3.5:1. DNA was added to 26μl DMEM Optimem (Invitrogen) and mixed with 1.9μl FuGene reagent. Complex formation was incubated 10-15min in RT before mixed with 40 000 ODInCas9 C86 cells seeded into a 24w. Plasmid sgRNAs targeted GFAP (GFAP-Cr1,2).

Synthetic two component sgRNAs in the form of crRNA and tracrRNA were purchased from IDT (Integrated DNA Technologies, Coralville, IA) (Supp. Info Table 2). Preparation of synthetic sgRNAs were performed according to manufacturer’s instructions by preparing 100μM stock concentrations of crRNA and tracrRNA, respectively, before mixing crRNA, tracrRNA, and Duplex Buffer (1:1:10). RNA mixture was heated for 5min at 95 degrees followed by cooling to room temperature. RNA was delivered using either lipid-based transfection (RNAiMAX) or electroporation. RNAiMAX (Invitrogen) and RNA was first mixed with DMEM Optimem (Invitrogen), separately, before combining the two solutions followed by a 20min incubation. ODInCas9 C86 cells were mixed with the lipid complex solution before seeding. For single sgRNA targeting (PIGM-Cr1,2) 30pmol RNA mixture was transfected per 40000 cells while for dual sgRNA targeting (PIGM-Cr1&2, GFAP-Cr1&2) 15pmol RNA mixture per target was transfected per 40000 cells. RNA delivery using the NEON electroporation system (ThermoFisher) was performed by applying 1600v, 10ms and 3 pulses of OVARC8 ODInCas9 pool mixed with RNA, 50,000 cells and 1 μL of sgRNA (MCT-Cr1&2) per 10 μL tip.

*In-vitro* transcribed sgRNA (GFAP-Cr1&2) was purchased from Eupheria Biotech (Dresden, Germany) and loaded into lipid nano particles (LNPs). Different amounts of LNPs (100ng-4000ng) were mixed with 40 000 ODInCas9 C86 cells and seeded into a 24w.

#### Proliferation Imaging

Human iPSC ODInCas9 line C86 was either non-induced or pulse induced (1h with 10μg/ml Dox following washout) 6h before synthetic sgRNA transfection using GFAP and Alu targets (Alu-Cr1,2, GFAP-Cr1&2). Growth curves were measured using IncuCyte Zoom microscope (Essen Bioscience Inc., Ann Arbor, MI) for 42h post transfection. OVCAR8 ODInCas9 cells were induced 24h before transfection and kept at 100 ng/mL Dox during the experiment. Transfection of synthetic spCas9 sgRNAs was done according to standard protocol targeting MCT1 (MCT1-Cr1&2) or CDK12 (CDK12-Cr1&2) in dual combination, respectively. Alu targets (Alu-cr1,2) were used as viability control. All targets were performed in triplicates.

#### FLAER Assay

FLAER assay (Cedarlane, Canada) tags mammalian GPI anchors through an AlexaFluor488 labeled version of aerolysin. Human iPSC ODInCas9 line C86 was pulse induced 1h with 10μg/ml Dox followed by washout 6h before standard synthetic sgRNA transfection targeting 3 sites of the *PIGM* gene (PIGM-Cr1,2,3) in single and dual-combination. ODInCas9 system was left to turn off for 7 days post transfection before assessment using FACS instrument BD LSRFortessa (BD Bioscience) and confocal microscopy instrument Cell Voyager 7000S (CV7000S, Yokogawa, Japan). Cells were detached using TryPLE (Invitrogen) and washed before cell suspension was incubated at room temperature for 15min at a final FLAER concentration of 0.5μM in 100μl volume. Cells were washed twice and resuspended in 100μl DPBS containing 2% fetal bovine serum (both from Invitrogen) before 5μl of was used to plate cells for microscopy imaging and the rest used for FACS analysis. Data were subsequently analyzed using FlowJo to assess reduction of GFP expression.

#### On Off activation of ODInCas9 transgene

Validation of on off capacity of the ODInCas9 transgene hiPSC ODInCas9 C86 was either non-induced or pulse induced 1h with 10μg/ml Dox followed by washout 6h before transfected with dual synthetic sgRNA transfection targeting *GFAP* (GFAP-Cr1&2) at d0. Induced cells at d0 was divided at d4 into non-induced or 2^nd^ pulse induced (1h with 10μg/ml Dox followed by washout) cells before transfected with dual synthetic sgRNA transfection targeting *MYLIP* (MYLIP-Cr1&2). Non induced cells at d0 where also transfected with dual synthetic sgRNA transfection targeting *MYLIP* and cultured to d7. DNA samples were collected at d4 and d7 for indel-assessment (see indel assessment method section). Protein collection was performed daily from d1-d8 to evaluate Cas9 protein expression (see western blot method section).

#### Whole transcriptome profiling by RNA sequencing and bioinformatics

Total RNA of 2-3 million cells was isolated using miRNeasy Kit according to the manufacturer’s instructions (Qiagen). The quality of the RNA was assessed by a standard sensitivity NGS fragment analysis kit on Fragment Analyzer (Advanced Analytical Technologies). All the samples had RNA integrity number >9.8 and were used for library preparation. One microgram of total RNA was used for each library. Illumina TruSeq Stranded Total RNA Ribo-Zero Human/Mouse/Rat Gold (Illumina) was used to construct ribosomal RNA depleted sequencing libraries. All libraries were quantified with the Fragment Analyzer, pooled in equimolar concentrations and quantified with Qubit Fluorometer (Invitrogen). Libraries sequenced >40M paired end reads using High Output Kit v2 (150 cycles) on an Illumina NextSeq500. Three biological replicates were sequenced per sample.

RNA-seq fastq files were processed using bcbio-nextgen (version 0.9.7) where reads were mapped to the genome with between 23.9 – 119.3 M mapped reads per sample (with a mean of 69.3 M). Gene level quantifications, counts and transcript per million (TPM), were generated with featurecounts (version 1.4.4) and sailfish (version 0.9.0), respectively, all within bcbio. All analyses were performed using R (version 3.4.0, https://www.r-project.org/). Differential gene expression were assessed with DESeq2 (version 1.14.1).

#### Indel/Edit detection

Gene editing was assessed by mismatch-specific endonuclease activity, either T7 endonuclease I (T7EI) or surveyor nuclease, on PCR amplified sequence of the target locus (Supp. Info Table 3). Genomic DNA was isolated using QuickExtract DNA Extraction Solution (Lucigen). PCR amplified target sequence was heated to 95°C and slowly (2C/s) cooled down to 25°C to form heteroduplexes before incubation with mismatch specific endonuclease which cleaves heteroduplexes enabling detection of indel formation. Cleaved products were analyzed by DNA electrophoresis using Novex TBE Gel 10% (ThermoFisher) or QIAxcel (Qiagen). Quantification of detected gene edit is performed as previously described [43].

#### Immunocytochemistry

24h after 100 ng/mL doxycycline treatment, the OVCAR8 ODInCas9 cells were fixed with methanol followed by permeabilization with 0.1% Triton X100. The samples were blocked in 10% goat serum and incubated overnight at 4°C with the mouse Cas9 antibody (7A9-3A3 Cell Signaling Technologies) 1:800 dilution in 1% BSA (Supp. Info Table 1). After washing, the cells were incubated with the goat anti-mouse DyLightTM 488 secondary antibody (ThermoFisher) 1:200 in 1% BSA. DAPI (0.5 ug/ml) was used to stain the nucleus of the cells. The stained cells were analyzed by the Image Xpress confocal microscope.

#### Generation of the OdinCas9 mouse

The target construct was electroporated into C57Bl/6N (Prx) ES cells. Neo-resistant clones were analyzed for correct integration into the R26 locus by PCR. Validation of transgene induction in mESC ODInCas9 cells was done by 48h Dox (10μg/ml) treatment assessed by GFP fluorescent signal and Cas9 protein expression. Three positive clones were further screened by Southern blot, using a Neo probe. One validated clone was expanded and injected into Balb/cAnNCrl blastocysts to generate chimeric mice. Chimeric C57Bl/6N OdinCas9 heterozygous males were bred to C57Bl/6NCrl females to generate experimental animals. Litters are genotyped using the following primers; ACGTTTCCGACTTGAGTTGC and GTGCAATCCATCTTGTTCA.

#### Animals

All mouse experiments were approved by the AstraZeneca internal committee for animal studies and the Gothenburg Ethics Committee for Experimental Animals (license numbers: 162–2015+ and 629-2017) compliant with EU directives on the protection of animals used for scientific purposes. Experimental heterozygous mice were generated by breeding male heterozygous ODinCas9 mice to female C57Bl/6NCrl mice (Charles River). Male and female ODinCas9 were bred to generate homozygous mice to understand the inheritance pattern of the cassette and to generate an experimental cohort. OdinCas9 mice were crossed to the described human knock in *PCSK9* mice [24, 44] to generate mice heterozygous for both constructs. In all experiments mice were randomised to groups to ensure that each group had similar body weight. Mice were housed in a temperature controlled room (21°C) with a 12:12 h light-dark cycle (dawn: 5.30 am, lights on: 6.00 am, dusk: 5.30pm, lights off: 6pm) and with controlled humidity (45–55%). Mice had access to a normal chow diet (R70, Lactamin AB, Stockholm, Sweden) and water *ad libitum*. Mice were checked daily and weighed weekly.

#### Guide RNA design and adeno-associated viral constructs

AAV (serotype 9) were custom generated by Vector Biolabs (Malvern, PA, USA). Each AAV consisted of 2 sgRNAs under a U6 promotor and contained a point mutation specific homology arm for each Kras G12D or G12C. The guide RNAs used for the non small cell lung cancer (NSCLC) studies have been validated previously (Supp Info Table 4) [13].

#### CRISPR-Cas9 induced editing with LNP or AAV in ODinCas9 mice

Induction of Cas9 required doxycycline hyclate (2mg/ml; Sigma Aldrich, MI, USA) supplemented with 1% sucrose added to the drinking water for 3 nights. To test the efficacy of guide delivery to the liver, OdinCas9 mice crossed with the human PCSK9 mice were injected intravenously with LNP containing guide for PCSK9 [24, 44], To generate models of NSCLC ODInCas9 mice (12-16 weeks), after 2 nights on Dox, mice were dosed by oropharyngeal aspiration with 1 × 10^10^ GC (*KrasG12D;Stk11^-/-^*), or 1 × 10^11^ GC (*KrasG12D;Trp53^-/-^, KrasG12C;Trp53^-/-^*) genome copies of adeno-associated (AAV9) diluted to a final volume of 50μl in PBS. 24h post dosing the mice were returned to normal drinking water.

#### NSCLC treatment studies

***KrasG12D;Trp53^-/-^*** ODInCas9 mice (16 weeks, male n=18, 33.5±2g) had 2 nights of dox were then dosed by oropharyngeal aspiration (OA) with 1 × 10^11^ GC of AAV9. Mice were weighed weekly and randomised to treatment group after 4 weeks ensuring each group had matched body weight. For 2 weeks, mice received Selumetinib (AZD6244, 25mg/kg, resuspended in 0.5% HPMC / 0.1% Tween 80) by oral gavage (BID, 8 hours apart). Once a week, 1 hour after Selumetinib dosing, Docetaxel (Taxotere, 15mg/kg, in physiological saline) was administered intravenously. ***KrasG12C;Trp53^-/-^*** ODInCas9 mice (10-12 weeks, male n=9, 28.7±2g and female n=13, 22.6±1g) had 2 nights dox and were OA dosed 1 × 10^11^ GC of AAV9. Mice were weighed weekly and randomized (3 male and 5 female) per treatment group after 12 weeks ensuring each treatment arm had matching body weight. For 4 weeks, mice were PO dosed (BID, 8 hours apart) with Selumetinib (AZD6244, 25mg/kg). Once a week, 1 hour after Selumetinib, Docetaxel (Taxotere, 15mg/kg) was administered intravenously.

#### Histology and immunohistochemistry

At necropsy, mice were euthanized under isoflurane anesthesia and lung, pancreas, liver, skeletal muscle, kidney, brain, gastrointestinal tract, reproductive system and spleen were collected in 4% neutral buffered formalin for assessment of Cas9 immunostained cell distribution. Further, lung tissue was inflated with 4% neutral buffered formalin and with other tissues immediately fixed in 4% neutral buffered formalin. Tissue was embedded in paraffin and prepared as 5μm thick sections. Sections were stained for hematoxylin and eosin for morphological characterization. To define the extracellular matrix, immune phenotype, pathway activation and the proliferative nature of the tumors, immunohistochemistry was performed with antibodies for αSMA, tenascin C, Nkx2.1, proSpC, CD31, F480, CD45R, CD4/8, pERK1/2, pMEK, Ki67, ⍰H2AX (supplier details and RRID# in Supp Info Table 1). Immunohistochemistry was performed using the Discovery Ultra (Ventana Medical Systems, Inc, AZ, USA). Antigen retrieval using cell condition fluid 1 was performed when required and the Discovery ChromoMap DAB was used as the detection system for all stains except CD4 Discovery Purple HRP and CD8 Discovery Teal HRP. All histological slides were blinded and examined using light microscopy (Carl Zeiss Microscopy GmbH, Jena, Germany) by an experienced board-certified pathologist. The tumor classification occurred as previously described and included atypical hyperplasia (grade 1), adenoma (grade 2), high grade adenocarcinoma (grade 3) and low grade adenocarcinoma (grade 4) [12].

#### NGS

Genomic DNA was extracted from whole lung tissue and isolated single adenocarcinomas from AAV injected mice using the Qiagen Puregene cell and tissue kit (Qiagen, Germany). The genomic DNA was amplified with adapter-containing gene-specific primers (refer to Supp Info Table 3), linker for forward primers: TCGTCGGCAGCGTCAGATGTGTATAAGAGACAG; the linker for reverse primers: GTCTCGTGGGCTCGGAGATGTGTATAAGAGACAG using Q5 Hot Start High Fidelity DNA polymerase (NEB) and sequenced by using a NextSeq500 Instrument (Illumina, California, United States). Read counts above 10K were achieved for the in vivo Trp53 and LNP delivery of guides. Read counts of 50K were achieved for all other analyzed alleles. Sequencing reads were demultiplexed using Illumina software and r1 and r2 FASTQ files were analyzed using CRISPResso [45]. Briefly, reads with a minimum average quality score of 30 were aligned to the reference sequence. A window of 30 base pairs centered on the predicted cleavage site was specified for the quantification of indels and base-editing outcome.

The frequencies of indel mutations were automatically calculated using CRISPResso. The frequency of mutated alleles was calculated based on the CRISPResso’s detected alleles, “Alleles_frequency_table.txt,” in NGS data as follows: first, all the detected alleles were imported into the Microsoft Excel program. Next, the number of reads for alleles with identical 30 nucleotides around the predicted cut site was consolidated. Alleles with a frequency of less than 0.01% were excluded from the analysis. Finally, the relative frequency of the 10 most frequent alleles among the consolidated alleles was calculated.

#### LNPs for in vitro experiments; Preparation, Characterization and Concentration

LNPs were manufactured using microfluidic mixing as described previously by Cullis [46, 47]. In brief, stocks of lipids were dissolved in ethanol to obtain a lipid concentration of 1-2 mM. The concentrations of the lipid stocks were carefully evaluated in order to maximize the number of loaded LNPs in order to increase the efficacy. xRNA (mRNA and/or gRNA) was diluted in RNase free 100 mM citrate buffer pH 3.0. The aqueous and ethanol solutions were mixed in a 3:1 volume ratio using a microfluidic apparatus NanoAssemblr (Precision NanoSystems Inc. Vancouver, Canada), at a mixing rate of 12 mL/min. LNPs were dialyzed overnight using Slide-A-Lyzer G2 dialysis cassettes (Thermo Scientific). The size of the LNPs was measured by dynamic light scattering (DLS) and performed on diluted samples in 10 mM phosphate buffer at 25°C. Data was collected at a scattering angle of 173° and the reported diameter is a mean of 3 values. Z-Average, number and volume weighted particle size distributions were calculated using a particle refractive index of 1.45 and an absorption of 0.001 and polydispersity (PDI) of the LNPs was determined using a Zetasizer Nano-ZS (Malvern Instruments Ltd). The size (Z-Average) of the LNPs prepared was in the range of 74-87 nm with a PDI of 0.14-0.21. The LNPs were concentrated to 100 μg/ml by ultra-spin filtration in Amicon Ultra 4, 30 000 kDa MWCO, centrifugal filter units and the size was re-measured by DLS in order to confirm the integrity of the LNPs. The encapsulation and concentration of mRNA were determined using the RiboGreen assay (Thermo Scientific) and the encapsulation efficiency for all samples was typically in the range of 92–96%.

#### Blood collection and triglyceride determination

Blood from vena saphena was collected in EDTA-coated tubes, centrifugated (7000g, 10min) and plasma was stored at −20°C. Plasma triglyceride was measured in all mice one week before treatment and 3 weeks after AAV transduction using an enzymatic colorimetric method (Cat. A11A01640; ABX Pentra 400; HORIBA Medical, Irvine, CA, USA).

#### SDS-Page and Western Blot analysis

Cells or tissue were lysed in RIPA buffer containing cOmplete Protease Inhibitor Cocktail (Roche, Germany) using a TissueLyser II (25Hzs for 1 min), cell debris was removed by centrifugation at 10000rpm for 10 min and protein concentration was determined using the Pierce™ BCA Protein Assay Kit (Thermo Fisher Scientific). Samples were prepared for SDS-PAGE (30μg loaded per well), separated by sodium dodecyl sulfate (SDS)-polyacrylamide gel electrophoresis (PAGE), transferred onto a PVDF membrane (Invitrogen, the Netherlands) and probed with rabbit pAb CRISPR/Cas9 (C15310258, Diagenode, Liège, Belgium, 1:5000), rabbit pAb GFP (ab290; Abcam, Cambridge, UK, 1:5000) and rabbit pAb GAPDH (9485; Abcam, Cambridge, UK, 1:10000) (Summarized in Supp. Table 1). Cell lysates were stained with MCT1 antibody (internal AZ generated 1:1000), rabbit pAb CDK12 antibody (Cell Signaling Technology, MA, USA, #11973, 1:1000), rabbit mAb β-Actin (Cell Signaling Technology, MA, USA D6A8) and rabbit mAb GAPDH (Cell Signaling Technology, 14C10, 1:3000). IRDye 800CW-labelled goat anti-rabbit (1:15000) and 680RD-labelled donkey anti-rabbit (1:150000) (LI-COR Biosciences, Cambridge, UK) were used for detection and blots were analyzed with the Odyssey infrared imaging system and software (LI-COR Biosciences) and quantified using the ImageJ software and the method of Thermo Fisher [48]. Quantification is performed by subtracting background signal from signal detected protein of interest (PI, Cas9/GFP) and normalization control (NC, Actin/GAPDH). A relative NC (rNC) value is derived by dividing each individual NC to the highest signal among the NC in each gel. Finally, all background subtracted PI intensity values are divided by each individually matching rNC.

#### BaseScope *in situ* hybridization assay

BaseScope assay from ACD was performed as described in Baker 2017 [25]. Briefly sections were baked, deparaffinized and dried. Pretreatment 1, 2 and then 3 were then conducted prior to BaseScope probe application (custom probes; I-BA-Mm-Kras-Donor-G12C-1zz-st, I-BA-Mm-Kras-Donor-G12D-1zz-st). The slides were then incubated in reagents AMP0, AMP1, AMP2, AMP3, AMP4, AMP5 and AMP6 with rinsing between each step. Finally, slides were incubated with Fast Red and then counterstained with hematoxylin mounting in VectaMount (Vector labs, Burlingame, CA). Semi-quantitative assessment of the BaseScope staining was performed using the ACD guidelines.

#### MRI

All MRI measurements were performed on a Biospec 9.4T/20 MRI scanner (Bruker BioSpin, Karlsruhe, Germany) equipped with a 400 mT/m actively shielded gradient system with ParaVision (PV5.1) software. For MRI measurement, a 50 mm i.d. quadrature resonator (m2m Imaging, Cleveland, Ohio) was used. Prior to the start of the MRI, animals were weighed and then anesthetized. Anesthesia was induced using 4% isoflurane (Attane vet^®^, 1000 mg/g, Piramal Healthcare, UK) in oxygen and maintained with 1.5–2.5% isoflurane in an air-oxygen mixture during the imaging session. The mouse was placed supine in a Plexiglas cradle and temperature maintained at ~36.5°C using a heating pad for the duration of the imaging session. MRI acquisitions were synchronized with the respiratory cycle using a respiratory pad placed under the abdomen of the animal to minimize physiological artefacts (SA Instruments, Stony Brook, NY).

The imaging protocol consisted of three scans. First, an ungated gradient echo localizer scan (TR/TE/α: 175 ms/3.8 ms/20°, number of averages (NA): 2, field of view (FOV): 30×30 mm, matrix size: 128×128) to localize the lungs. Second, a set of two gated multi-slice coronal scans acquired with a fat suppressed Rapid Acquisition with Relaxation Enhancement (RARE) pulse sequence with the following parameters: TR/TE/RARE factor: 826 ms/9 ms/8, NA: 8, FOV: 30×30 mm, in-plane resolution: 117 x 117 μm, number of slices: 9-10, inter-slice: 1.4 mm and slice thickness: 0.7 mm. The total acquisition time was approximately 10 mins per scan. Slice position of the second RARE scan were interleaved into the interslice gap of the first RARE scan in order to cover the whole thoracic cavity without any slice gaps.

##### Segmentation of tumor volume

The two multi-slice RARE scans were first imported into the software ImageJ^®^ (version 1.52a, NIH, MD, USA) and interleaved to obtain a single 3D image data set of the whole thoracic cavity (matrix size: 256×256×18-20). Tumor nodules were segmented semi-automatically using the image analysis software Analyze 12.0 (Biomedical Imaging Resource, Mayo Clinic, Rochester, MN). Segmentation was performed as follows: 1) lung regions comprising tumor nodules were manually contoured in each slice, 2) tumor nodules in each defined region were then segmented based on histogram thresholding. Volumes of tumor nodules (in mm^3^) were calculated by multiplying the number of segmented voxels by the voxel volume resolution. Total tumor volume was determined by summing the nodule volumes from each slice. To better evaluate and distinguish pulmonary nodules from normal lung parenchyma, MRI images are represented with inverted grayscale.

#### Statistical analysis

Data visualizations and statistical comparisons between groups were performed using GraphPad Prism (v7.02). ANOVA and when relevant, Dunnett’s post hoc test was performed for correction of multiple hypotheses. *P* < 0.05 was considered to be statistically significant. The level of significance in all graphs is represented as follows: * *P* < 0.05, ** *P* < 0.01, *** *P* < 0.001 and **** *P* < 0.0001. Exact *P* values are presented in the description of the results.

## ACKNOWLEDGEMENTS

We acknowledge Anne Goeppert and Ben Taylor for contributions in developing ODInCas9 cell lines. We thank Mikael Bjursell, Marie Johansson, Johan Johansson, Sara Torstensson, Liselotta Hallengren, Anna Thoren and Lillevi Kärrberg for help with the *in vivo* experiments. Thanks to Marianna Yanez Arteta who helped with the lipid nanoparticle formulation and Mike Firth for assistance with Amplicon Seq analysis.

## AUTHOR CONTRIBUTIONS

M.M., L.M.M. and M.B-Y. conceived the project; A.L., M.J.P., H.J., R.N., S.T.B, E.J.D, M.M. and M.B-Y. designed the study; X.X generated the transgene, A.L., H.J., J.B., generated cell lines; H.J., M.M. and M.B-Y. designed and generated the mouse model; A.L., M.J.P., H.J., R.N., J.B., M.C., and L.B. performed the experiments; A.L., M.J.P., F.S., A.B., B.A., S.T.B., C.P. M., E.J.D., M.B-Y. and M.M. contributed to data analysis and interpretation; F.S. and C.A.J., performed and reported on immunohistochemical studies; A.B., performed the MRI acquisition and data analysis; A.T.G. performed bioinformatics analysis; A.S. provided LNP formulations; A.L., M.J.P., S.T.B and M.M. wrote the manuscript, S.T.B., L.B., E.J.D, C.P.M., A.B., F.S. and M.M. contributed to manuscript revision and M.B-Y. and M.M. supervised the study. All authors discussed the results and approved the final manuscript.

## Figures

**Supplementary Figure 1 (associated to Figure 1).**
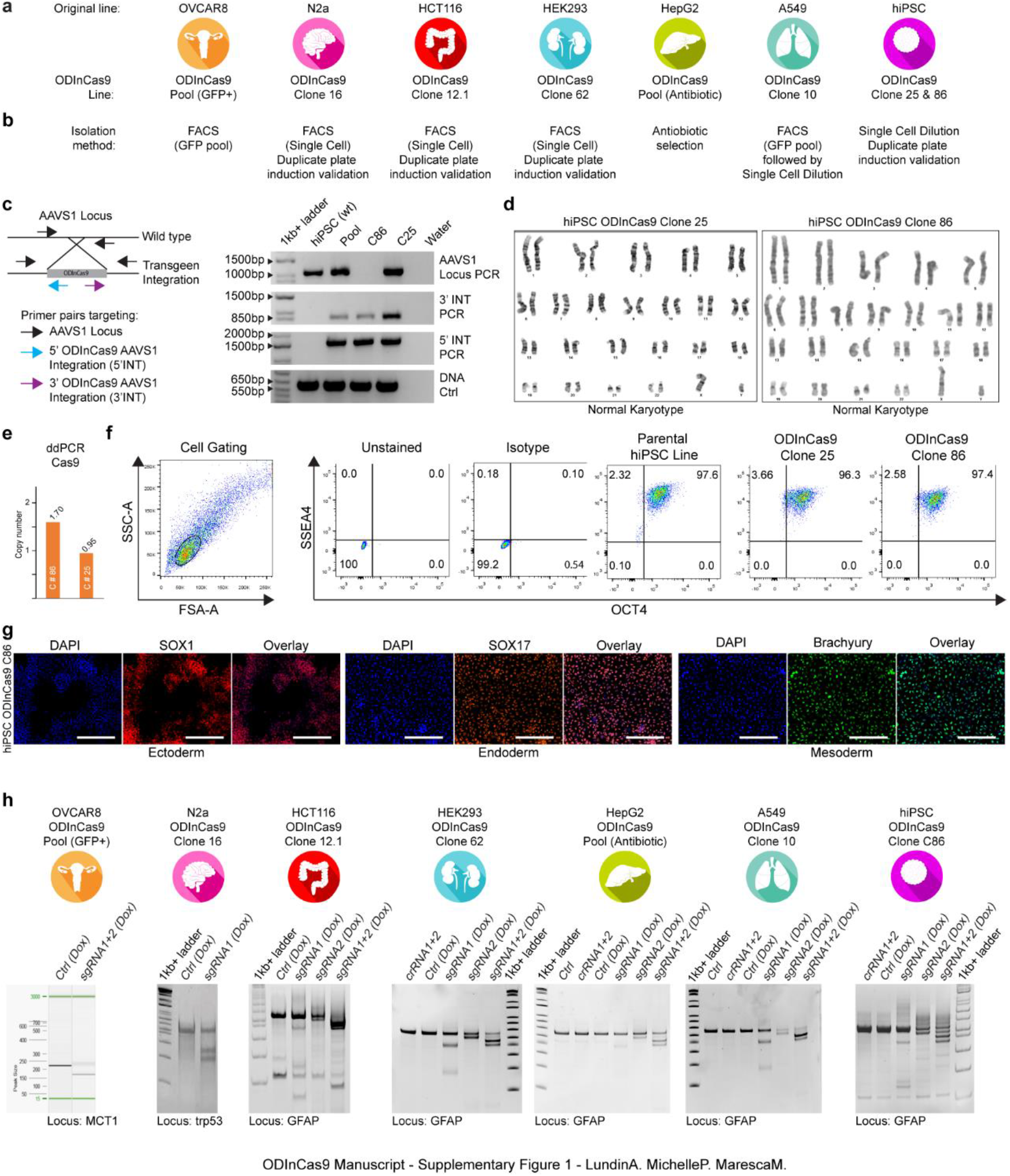
Cell line generation and validation. (**a**) ODInCas9 cell lines (**b**) isolated by various techniques including FACS of pool GFP+ cells (OVCAR8), single cell FACS with duplicate plate induction validation (N2a, HCT116, HEK293), antibiotic selection (HepG2) and single cell dilution with duplicate plate induction validation (hiPSC). (**c**) AAVS1 locus integrity and AAVS1 ODInCas9 transgene integration of hiPSC, hiPSC ODInCas9 pool, hiPSC ODInCas9 C25 and hiPSC OdinCas9 C86. (**d**) G-banding karyotype evaluation for hiPSC ODInCas9 C25 and C86. (**e**) Copy number integration of ODInCas9 identified by digital droplet PCR probing Cas9 sequence normalized to reference probe AP3B1 (**f**) FACS analysis of pluripotency markers SSEA4 and OCT4 of parental hiPSC line, hiPSC ODInCas9 C25 and C86. (**g**) Differentiation of hiPSC ODInCas9 C86 into the three germ layers assessed by immunocytochemical staining of markers for ectoderm (SOX1), endoderm (SOX17) and mesoderm (TBXT). (**h**) ODInCas9 induction and nuclease activity assessed 48h post transfection by mismatch endonuclease activity assay targeting; MCT1 in OVCAR8 by paired sgRNA, trp53 in N2A by single sgRNA, GFAP in HCT116, HEK293, HepG2, A549 and hiPSC by single sgRNA and paired sgRNAs. Dox = Doxycycline induced cell cultures. Scale bar: 300μm

**Supplementary Figure 2 (associated to Figure 2).**
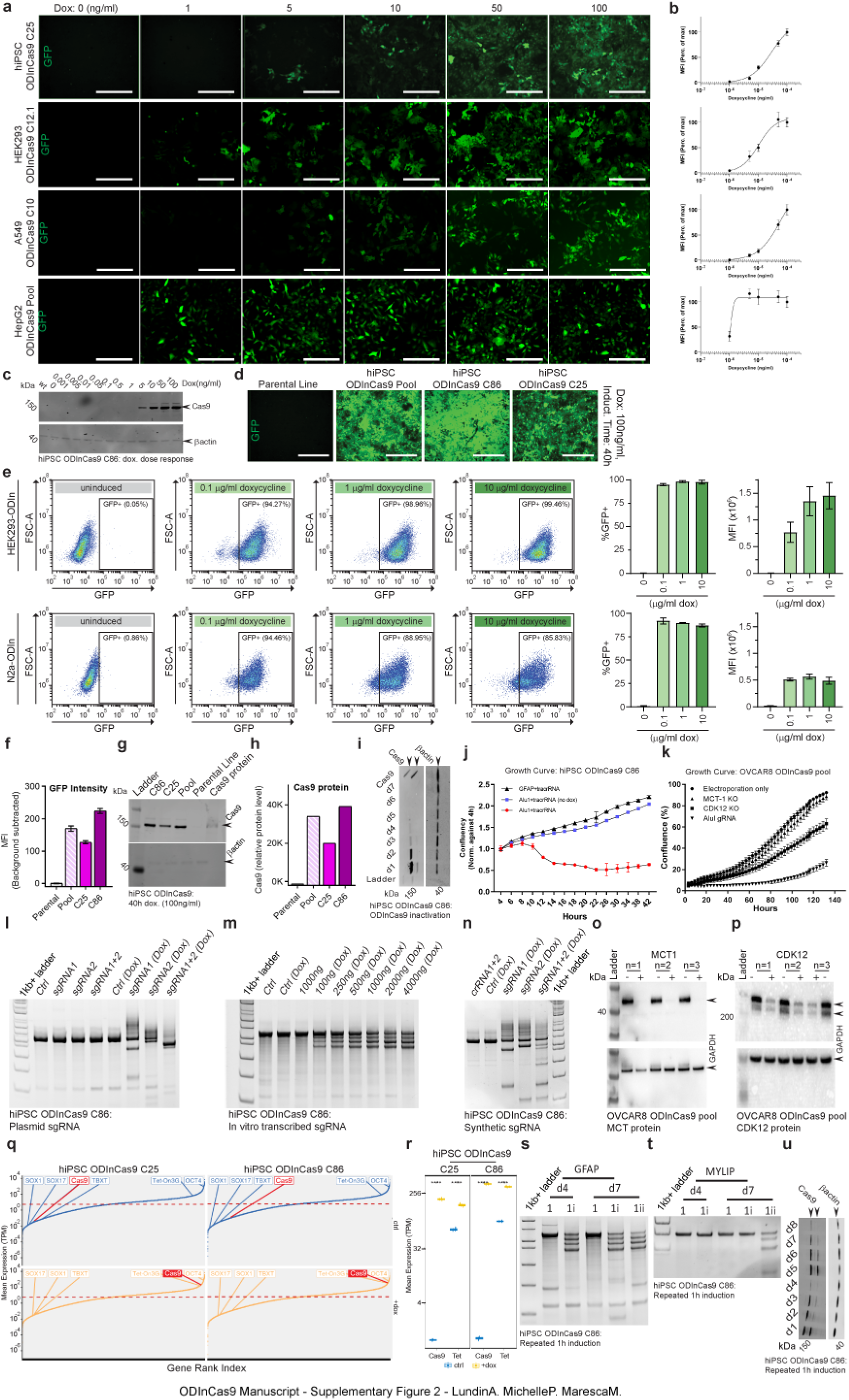
Regulation and application of ODInCas9 system. Transgene activation of ODInCas9 lines assessed by (**a**) GFP expression and (**b**) MFI (imaging, n=9 per concentration) normalized to maximum fluorescence intensity at highest dox concentration). (**c**) Cas9 protein expression of ODInCas9.C86 treated with dox concentration 0-100ng/ml. (**d**) GFP expression of hiPSC parental line, ODInCas9.Pool, ODInCas9.C86/C25 treated with 100ng/ml for 40h. (**e**) Fluorescent analysis of HEK293.C12.1 and N2A.C.16. 48h post dox treatment (0.1, 1 and 10μ/ml) (**f**) GFP MFI of hiPSC parental line, ODInCas9.Pool, ODInCas9.C86/C25 treated with 100ng/ml for 40h (imaging, n=9 per concentration; background subtracted). (**g**) Cas9 protein expression and (**h**) quantification of hiPSC parental line, ODInCas9.Pool, ODInCas9.C86/C25 treated with 100ng/ml for 40h (normalized to actin concentration). (**i**) Time course study of Cas9 protein expression following 1h of 10μg/ml dox induction. (**j**) Cell confluency normalized against earliest readout of seeding density at 4h. Transfection at 0h of sgRNA targeting GFAP locus (control) and Alu sites with and w/o dox (100ng/ml). (**k**) Cell confluency following electroporation at 0h of sgRNA targeting MCT1, CDK12 and Alu sites. Mismatch endonuclease activity assay of GFAP locus PCR product targeted by sgRNA and paired sgRNA delivered as (**l**) plasmid sgRNA using liposomal vehicle, (**m**) in vitro transcribed sgRNA using lipid nanoparticles at increasing concentrations and (**n**) synthetic sgRNA using liposomal vehicle. (**o**) MCT1 and (**p**) CDK12 protein expression following electroporation at 0h of sgRNA targeting MCT1 and CDK12 of activated and non-activated OVCAR8.ODInCas9.Pool cells. (**q**) Expression of *Cas9* and *Tet-On3G* transcript level relative to whole transcriptome in relation to biomarkers *SOX1, SOX17, TBXT* and *OCT4* of hiPSC.ODInCas9.C25/C86 in the absence or 48h treatment with dox (10μg/ml). Genes without expression in any replicate were filtered out. Only protein coding genes are shown. Mean expression of genes across the 3 replicate is shown on the y axis. (**r**) *Cas9* and *Tet-On3G* transcript levels (TPM) of hiPSC.ODInCas9.C25/C86 in the absence or 48h treatment with dox (10μg/ml). Mismatch endonuclease activity assay of (**s**) GFAP and (**t**) MYLIP locus PCR product targeted by paired sgRNA. Transfection of sgRNAs targeting GFAP and MYLIP performed at d0 and d4, respectively, and genomic DNA isolated at d4 and d7. (**u**) Time course study of Cas9 protein expression during repeated activation of ODInCas9 system by 10μg/ml dox 1h inductions at d0 and d4. Data shown as mean ± SEM, MFI = mean fluorescent intensity, TPM = transcripts per million. **** = <1e-4. Scale bar: 300μm

**Supplementary Figure 3 (associated to Figure 3).**
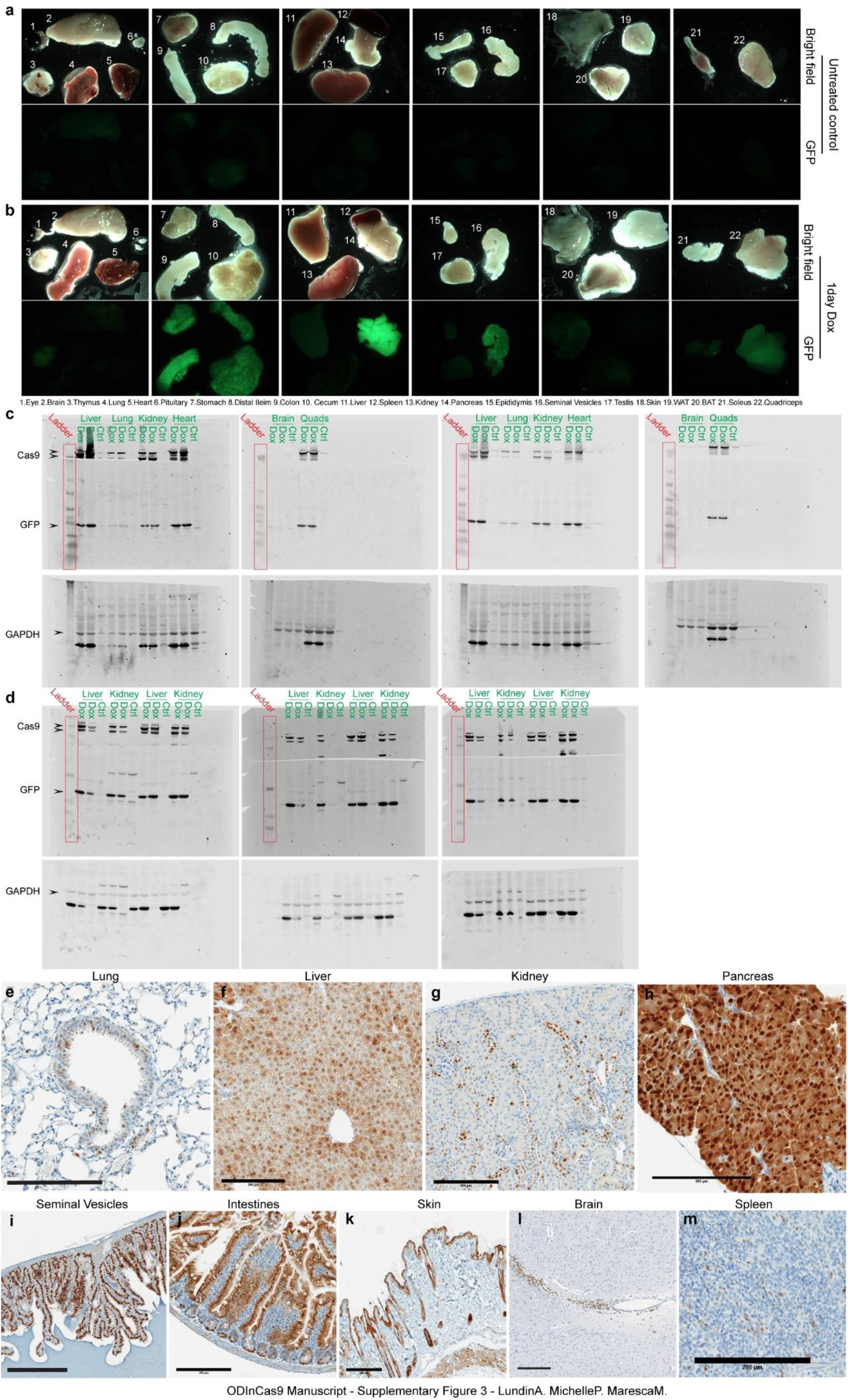
Regulation of the ODInCas9 system *in vivo*. Evaluation of organ GFP intensity of (**a**) untreated control (**b**) 1day dox stimulation. (**c**) Western blots of Cas9, GFP and GAPDH for liver, lung, kidney, heart, brain, quadriceps of heterozygote ODInCas9 mice without or with 3-day doxycycline stimulation. (**d**) Western blots of Cas9, GFP and GAPDH for liver and kidney of homozygous, heterozygous ODInCas9 mice and wild type mice after 3-day doxycycline stimulation. Immunohistochemistry section of (**e**) lung, (**f**) liver, (**g**) kidney, (**h**) pancreas, (**i**) seminal vesicles, (**j**) intestines, (**k**) skin, (**l**) brain and (**m**) spleen staining for Cas9 of heterozygote ODInCas9 mice.

**Supplementary Figure 4A (associated to Figure 4).**
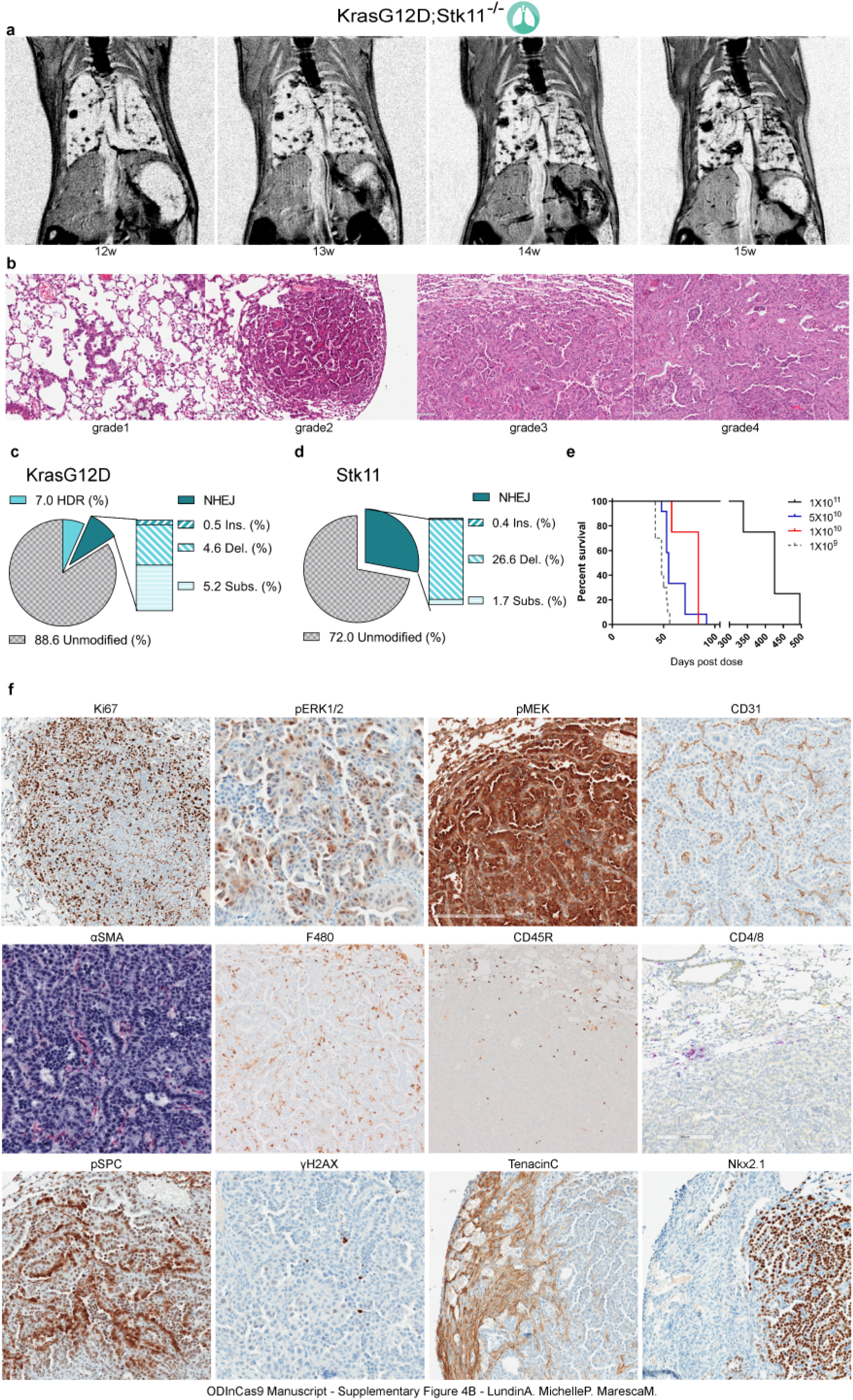
Modelling NSCLC in the ODInCas9 *KrasG12D;Stk11^-/-^* model. (**a**) Time course of tumor development by MRI following the same mice (**b**) Representative HE staining of mouse lung capturing the various stages of lung adenocarcinoma (**c,d**) Classification of amplicon sequencing of lung cancer tissue at *Kras* and *Stk11* locus (**e**) Kaplan-Meier survival curve for 4 different AAV9 viral titers (GC) (**f**) Immunohistochemistry section of lung tumors staining for Ki67, pERK, pMEK, CD31, αSMA, F480, CD45R, CD4/8, pSPC, ɣH2AX, Tenascin C and Nkx2.1.

**Supplementary Figure 4B (associated to Figure 4).**
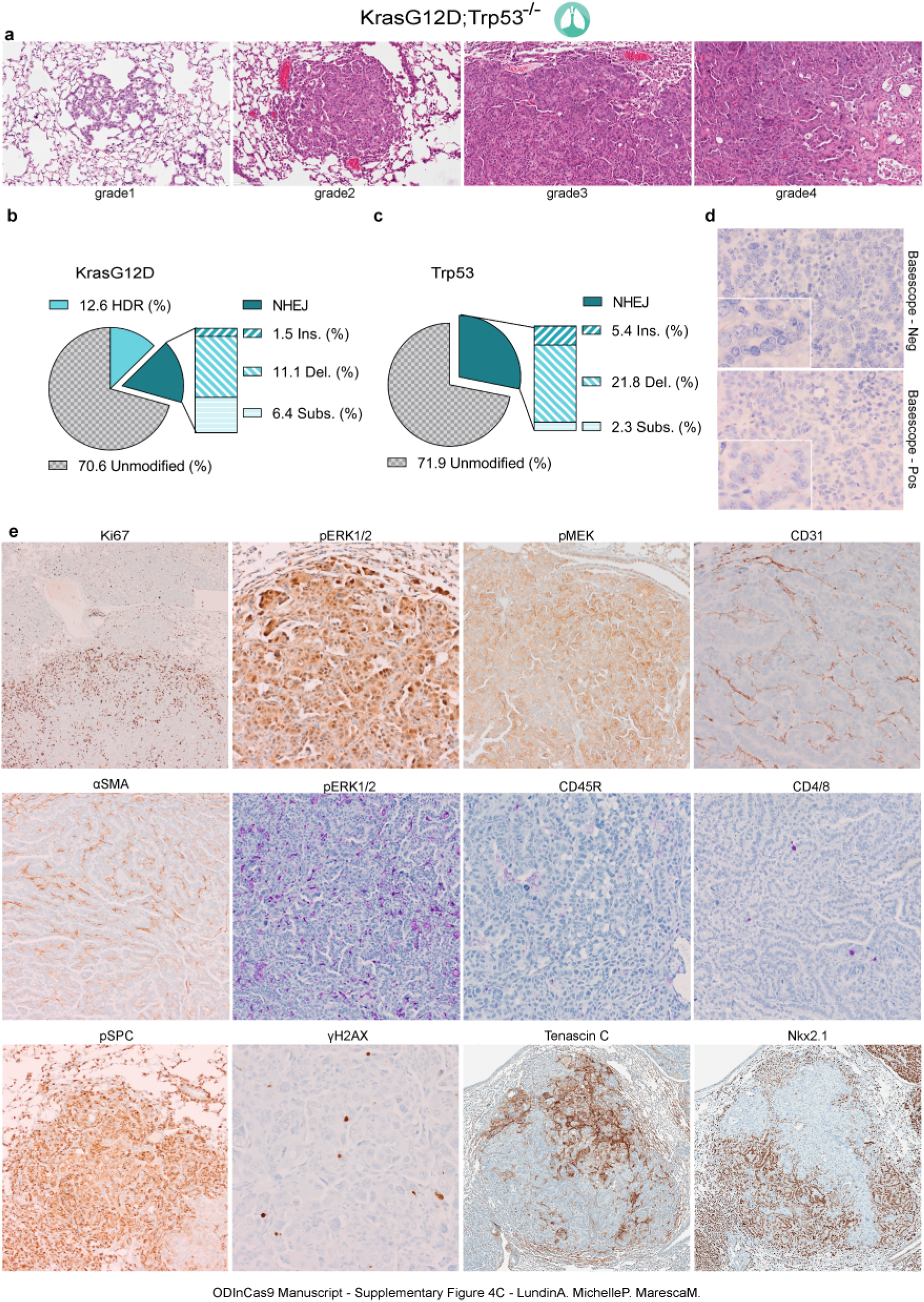
ODInCas9 *KrasG12C;Trp53^-/-^* NSCLC model. (**a**) Representative HE staining of mouse lung capturing the various stages of lung adenocarcinoma (**b,c**) Classification of amplicon sequencing of lung cancer tissue at *Kras* and *Trp53* locus (**d**) DNA *in situ* hybridization targeting KrasG12C (**e**) Immunohistochemistry staining of lung tumors for Ki67, pERK, pMEK, CD31, αSMA, F480, CD45R, CD4/8, pSPC, ɣH2AX, Tenascin C and Nkx2.1

**Supplementary Figure 5 (associated to Figure 5).**
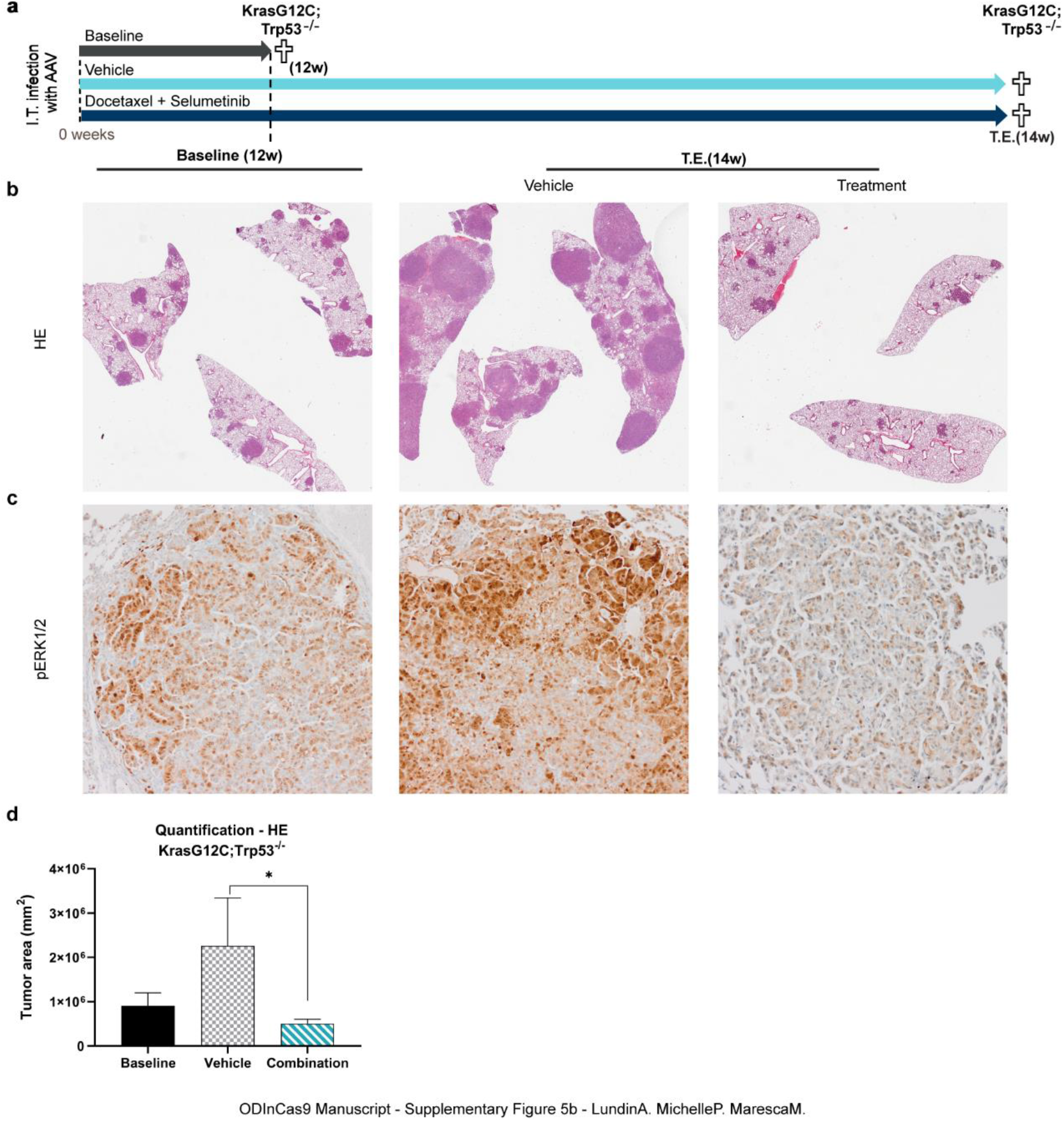
Pre-clinical efficacy study of ODInCas9 *KrasG12C;Trp53^-/-^* NSCLC model. **(a)** Timeline for treatment study. ODInCas9 mice (n=22) were induced with dox, dosed with AAV and allocated to treatment arm at 12w when significant tumor burden is observed. Mice were treated for 2-weeks with either vehicle or Docetaxel + Selumetinib combination treatment. Evaluation at baseline and treatment end (T.E) by (**b**) lung tissue histology (HE) (**c**) pERK1/2 immunohistochemical staining. (**d**) Quantification of tumor burden at baseline, vehicle and combinatory treatment using histology (ANOVA, F (2,9)= 6.490, p=0.0180).

