## Supplemental table 1 for "ObLiGaRe doxycycline Inducible (ODIn) Cas9 system driving pre-clinical drug discovery, from design to cancer treatment"

Supplementary Table 1

| **Table 1 -** |  |  |  |  |  |
| --- | --- | --- | --- | --- | --- |
| Reagents for immunocytochemistry, immunohistochemistry, Western blotting and BaseScope | | | |  |  |
| Research Resource Identification Portal (RRIDs) - https://scicrunch.org/resources | | | |  |  |
| ICC |  |  |  |  |  |
| **Antibody** | **Vendor** | **Catalog number** | **clone** | **RRID Portal #** | **Figure** |
| Cas9 | Cell Signaling Technology, MA, USA | 7A9-3A3 | monoclonal | AB_2750916 | 2c, Supp 2c |
| beta actin | Cell Signaling Technology, MA, USA | 8457 | monoclonal | AB_10950489 | 2c, Supp 2c |
| anti mouse IgG (H+L) DyLight 488 | ThermoFisher Scientific, MA, USA | 35502 | polyclonal | AB_844397 | 2c, Supp 2c |
| IHC |  |  |  |  |  |
| **Antibody** | **Vendor** | **Catalog number** | **clone** | **RRID Portal #** | **Figure** |
| Cas9 | Diagenode, Liège, Belgium | C15310258-20 | polyclonal | AB_2715516 | Supp 3e-m |
| alpha SMA | Cell Signaling Technology, MA, USA | 19245 | D4K9N | AB_2734735 | 4f, Supp 4A f, Supp 4B f |
| CD31 | Abcam, Cambridge, UK | ab28364 | polyclonal | AB_726362 | 4f, Supp 4A f, Supp 4B f |
| F4/80 | Cell Signaling Technology, MA, USA | 70076 | D2S9R | AB_2799771 | 4f, Supp 4A f, Supp 4B f |
| CD45R | BD Pharmingen | 553084 | RA3-6B2 | AB_394614 | 4f, Supp 4A f, Supp 4B f |
| CD4 | Abcam, Cambridge, UK | ab183685 | EPR19514 | AB_2686917 | 4f, Supp 4A f, Supp 4B f |
| CD8 | Cell Signaling Technology, MA, USA | 98941 | D4W2Z | AB_2756376 | 4f, Supp 4A f, Supp 4B f |
| pSPC (prosurfactant prot C) | Abcam, Cambridge, UK | ab90716 | polyclonal | AB_10674024 | 4f, Supp 4A f, Supp 4B f |
| gH2AX(phospho-Histone) | Cell Signaling Technology, MA, USA | 2577 | Ser139 | AB_2118010 | 4f, Supp 4A f, Supp 4B f |
| Tenascin C | Millipore, MA USA | AB19011 | polyclonal | AB_2203804 | 4f, Supp 4A f, Supp 4B f |
| NKx2.1 (TFF1) | Abcam, Cambridge, UK | ab76013 | EP1584Y | AB_1310784 | 4f, Supp 4A f, Supp 4B f |
| MAC2 | Cedarlane | CL8942AP | M3/38 | AB_10060357 | 4f, Supp 4A f, Supp 4B f |
| Ki67 | Abcam, Cambridge, UK | ab15580 | polyclonal | AB_443209 | 4f, Supp 4A f, Supp 4B f |
| pERK 1/2 (phospho-p42/44) | Cell Signaling Technology, MA, USA | 4376 | (Thr202/Tyr204) (20G11) | AB_331772 | 4f, Supp 4A f, Supp 4B f, 5d, Supp 5A d |
| pMEK (Phospho-MEK1/2) | Cell Signaling Technology, MA, USA | 2338 | (Ser221) (166F8) | AB_490903 | 4f, Supp 4A f, Supp 4B f |
| Western blot |  |  |  |  |  |
| **Antibody** | **Vendor** | **Catalog number** | **clone** | **RRID Portal #** | **Figure** |
| CRISPR/Cas9 | Diagenode, Liège, Belgium | C15310258 | polyclonal | AB_2715516 | 3f, Supp 3c, d |
| GFP | Abcam, Cambridge, UK | ab290 | polyclonal | AB_303395 | 3g, Supp 3c, d |
| GAPDH | Abcam, Cambridge, UK | 9485 | polyclonal | AB_307275 | 3f,g Supp 3c,d |
| MCT1 antibody | internally manufactured AstraZencea |  |  |  | Supp 2o |
| CDK12 | Cell Signaling Technology, MA, USA | 11973 | polyclonal | AB_2715688 | Supp 2p |
| GAPDH | Cell Signaling Technology, MA, USA | 2118 | 14C10 | AB_561053 | Supp 2o, p |
| goat anti-rabbit 800CW | LI-COR Biosciences, Cambridge, UK | 925-32211 | polyclonal | AB_2651127 | 3g, Supp 3c, d |
| donkey anti-rabbit 680RD | LI-COR Biosciences, Cambridge, UK | 925-68073 | polyclonal | AB_2716687 | 3f,g Supp 3c,d |
| BaseScope |  |  |  |  |  |
| **Probe set** | **Probe name** | **catalog number** | **Target** |  |  |
| Kras G12D | I-BA-Mm-Kras-Donor-G12D-1zz-st | 720508 | Mus musculus v-Ki-ras2 Kirsten rat sarcoma viral oncogene homolog (KRAS), donor KrasG12D, mRNA |  |  |
| Kras G12C | I-BA-Mm-Kras-Donor-G12C-1zz-st | 720518 | Mus musculus v-Ki-ras2 Kirsten rat sarcoma viral oncogene homolog (KRAS), donor KrasG12C, mRNA |  |  |
