## Supplemental table 2 for "ObLiGaRe doxycycline Inducible (ODIn) Cas9 system driving pre-clinical drug discovery, from design to cancer treatment"

Supplementary Table 2

| **Table 2 -** |  |  |  |
| --- | --- | --- | --- |
| CrRNA sequences for genomic targets in human and mice. | | |  |
| **Guide RNA** | **Target** | **crRNA Seqeunce** | **Figure** |
| Cr1-GFAP | Human GFAP | GGGTGCCAGGACCCAGACGG | 1f, Supp 1h, 2j-k, 2o-p, Supp 2l-n, Supp 2s |
| Cr2-GFAP | Human GFAP | GGAGACCCGGGCCAGTGAGC | 1f, Supp 1h, 2j-k, 2o-p, Supp 2l-n, Supp 2s |
| Cr1-MYLIP | Human MYLIP | AGGGCAGAAACTGCTCAT | 2q, Supp 2t |
| Cr2-MYLIP | Human MYLIP | TGGAAAACTATGGCATAGAA | 2q, Supp 2t |
| Cr1-PIGM | Human PIGM | AACTTGAAGGTGGCTCCAGC | 2l-n, Supp 2n |
| Cr2-PIGM | Human PIGM | TGTCCGTATACCTCACGTGC | 2l-n, Supp 2n |
| Cr3-PIGM | Human PIGM | TGGATACTGCCATAGGCAGG | 2n |
| Cr1-p53 | Mouse Trp53 | GTGTAATAGCTCCTGCATGG | 1f, Supp 1h |
| Cr1-MCT1 | Human MCT1 | CGTATAGTCATGATTGTTGG | 1f, Supp 1h, Supp 2k, Supp 2o |
| Cr2-MCT1 | Human MCT1 | ACAGACGTATAGTTGCTGTA | 1f, Supp 1h, Supp 2k, Supp 2o |
| Cr1-CDK12 | Human CDK12 | GTCGGTCAGTCCCCCTTACA | Supp 2k, Supp 2p |
| Cr2-CDK12 | Human CDK12 | GACGACCGTCTCCTGCTATA | Supp 2k, Supp 2p |
| Cr-Alu1 | Alu | CCTGTAATCCCAGCA | Supp 2j-k |
| Cr-Alu2 | Alu | CACTTTGGGAGGCCG | Supp 2j-k |
