## Supplemental table 3 for "ObLiGaRe doxycycline Inducible (ODIn) Cas9 system driving pre-clinical drug discovery, from design to cancer treatment"

Supplementary Table 3

| Table 3 - | | |  |  | |  |  | |  | |
| --- | --- | --- | --- | --- | --- | --- | --- | --- | --- | --- |
| Primers and probes used in this study | | | | | | |  | |  | |
| **Primer target** | | **Species** | | | **Function** | **FwPrimer** | | **RevPrimer** | | **Figure** |
| GFAP | | Human | | | Cel1 | TCATCATGGTCCAACCAACC | | GAAGCGAACCTTCTCGATGT | | 1f, Supp 1h, 2j-k, 2o-p, Supp 2l-n, Supp 2s |
| MYLIP | | Human | | | Cel1 | CTCTTGGTGTCCTCCAGCAT | | TTGGGCTAAAGTCATCTTTACAA | | 2q, Supp 2t |
| PIGM | | Human | | | Cel1 | AGTCTGCAGTCGTTTCGGTT | | TCTATGGTTTCGCGGTGCAT | | 2l, Supp 2n |
| MCT1 | | Human | | | Cel1 | TGTGAGGGAGCAGTTTCCTT | | GCCAGCCATAAACTAATGCTTC | | 1f, Supp 1h |
| AAVS1 | | Human | | | 3’INT | TGGCGTTACTATGGGAACATACG | | AAAGAGTCCCCAGTGCTATCTGG | | Supp 1c |
| AAVS1 | | Human | | | 5’INT | GTTTTTCTGGACAACCCCAAAGT | | GCAATATCACGGGTAGCCAA | | Supp 1c |
| AAVS1 | | Human | | | WT | GTTTTTCTGGACAACCCCAAAGT | | AAAGAGTCCCCAGTGCTATCTGG | | Supp 1c |
| PCSK9 | | Human | | | NGS | CCTGCAGGCCCAGGCTGC | | TCCGCCCGGTACCGTGGA | | 3l |
| Trp53 | | Mouse | | | Cel1 | TAACAGCAGTCTCTGGGAGAAG | | CTAAGCCCAAGAGGAAACAGAG | | 1f, Supp 1h |
| Kras | | Mouse | | | NGS | AGGCCTGCTGAAAATGACTGAGTA | | AAGCACGGATGGCATCTTGGACC | | 4c, Supp 4Ac, Supp 4B b |
| Trp53 | | Mouse | | | NGS | TGTAGTGAGGTAGGGAGCGAC | | GAACCAAAGAGCGTTGGGC | | 4d, Supp 4B c |
| Stk11 | | Mouse | | | NGS | GACACCTTCATCCACCGCA | | TGGCACCTACTTCTTGACGTT | | Supp 4B b |
