## Supplemental table 4 for "ObLiGaRe doxycycline Inducible (ODIn) Cas9 system driving pre-clinical drug discovery, from design to cancer treatment"

Supplementary Table 4

| Table 4 - | |  |  |  |  |
| --- | --- | --- | --- | --- | --- |
| guide RNAs used in this study | | | |  |  |
| **Primer target** | **Species** | | **Method** | **Sequence sgRNA** | **Figure** |
| PCSK9 | Human | | LNP delivery | CAGGTTCCACGGGATGCTCT | 3l, m |
| Kras | Mouse | | AAV | GCAGCGTTACCTCTATCGTA | 4, Supp 4A, Supp 4B, 5, Supp 5A |
| Trp53 | Mouse | | AAV | GTGTAATAGCTCCTGCATGG | 3l, 4, Supp 4B, 5, Supp 5A |
| Stk11 | Mouse | | AAV | ACTCCGAGACCTTATGCCGC | Supp 4A |
